# *Plasmodium falciparum* stomatin-like protein forms a putative complex with a metalloprotease in distinct mitochondrial loci

**DOI:** 10.1101/2024.07.18.604071

**Authors:** Julie M.J. Verhoef, Ezra T. Bekkering, Cas Boshoven, Megan Hannon, Felix Evers, Nicholas I. Proellochs, Cornelia G. Spruijt, Taco W.A. Kooij

**Affiliations:** Department of Medical Microbiology, Radboud University Medical Center, Nijmegen, The Netherlands; Department of Molecular Biology, Faculty of Science, Radboud University Nijmegen, Nijmegen, The Netherlands

## Abstract

Members of the <u>S</u>tomatin, <u>P</u>rohibitin, <u>F</u>lotillin and <u>H</u>flK/C (SPFH) protein family form large membrane anchored or spanning complexes and are involved in various functions in different organelles. The human malaria causing parasite *Plasmodium falciparum* harbors four SPFH proteins, including prohibitin 1 and 2, prohibitin-like protein (PHBL), and stomatin-like protein (STOML), which all localize to the parasite mitochondrion. In the murine model parasite *Plasmodium berghei, STOML* appears essential for asexual blood-stage (ABS) development and is localized to puncta on mitochondrial branching points in oocyst stages. In this study, we show that deletion of *STOML* causes a significant growth defect and slower ABS development, while sexual-stage development remains unaffected. Parasites lacking *STOML* were not more sensitive to respiratory chain targeting drugs, rendering a function of STOML in respiratory chain assembly unlikely. Epitope tagging of endogenous STOML revealed a distinct punctate localization on branching points and endings of the ABS mitochondrial network. STOML resides in a large protein complex and pulldown experiments identified a zinc dependent metalloprotease, FtsH, as a likely interaction partner. The predicted AlphaFold2 structure of STOML shows high similarity with the bacterial HflK/C, which has been shown to form a large vault-like structure around bacterial FtsH hexamers. Combined, our results suggest that a similar STOML-FtsH complex localized to specific loci of *P. falciparum* mitochondria facilitate the parasite’s ABS development.

## Introduction

Malaria is an infectious disease caused by *Plasmodium* parasites, which takes more than 600,000 mostly young lives annually^1^. *Plasmodium falciparum* is the most virulent malaria causing species. Resistance to current antimalarial drugs is spreading fast, emphasizing the need for the continuous development of novel antimalarial compounds. *Plasmodium* parasites harbor a unique mitochondrion that differs considerably from human mitochondria, which makes it a suitable drug target of antimalarial compounds such as atovaquone, DSM265, ELQ300, and proguanil^2,3^.

The mitochondrion consists of an inner and outer membrane, which are both rich in large protein complexes. Indeed, the inner mitochondrial membrane (IMM) is considered one of the most protein-rich membranes in any cell-type and contains large multiprotein complexes, such as respiratory chain complexes, ATP synthase, and the mitochondrial contact site and cristae organizing system (MICOS). Similarly to all other biological membranes, the mitochondrial membranes are organized into domains of distinct protein and lipid composition^4,5^. These membrane microdomains are important for the spatial and temporal control of membrane protein complex assembly and regulation^4^. SPFH (<u>S</u>tomatin, <u>P</u>rohibitin, <u>F</u>lotillin and <u>H</u>flK/C) family proteins are enriched in eukaryotic and prokaryotic membrane microdomains of various organelles, such as plasma membrane, nucleus, endoplasmic reticulum (ER), and mitochondria^6^. The common feature of SPFH proteins is the presence of the highly conserved SPFH or Band-7 protein domain^7^. These proteins form large self-oligomerizing membrane-spanning or membrane-anchored complexes and have been indicated in a diverse set of functions^6^. In human mitochondria, a subset of SPFH proteins, including two prohibitins (PHB1 and PHB2) and stomatin-like protein 2 (SLP2), localizes to the IMM. PHB1 and PHB2 form a large protein complex together, which has been indicated to play a role in mitochondrial protein degradation, cristae formation, mitochondrial dynamics, cell cycle regulation, and apoptosis^8–11^. SLP2 localizes to cardiolipin enriched membrane microdomains, where it interacts with and controls stability of the PHB complex^12,13^. The PHB and SLP2 complexes are both important for the formation and stability of the respiratory chain complex and mitochondrial translation^8,13–17^. They reside in large supercomplexes with metalloproteases and assert their proteolytic function through regulation of metalloprotease activity, similar to their bacterial family member HflIK/C^8,11,12,18–21^.

*Plasmodium* parasites harbor three conserved SPFH proteins: PHB1, PHB2, and stomatin-like protein (STOML), as well as an unusual prohibitin-like protein (PHBL). PHBL is specific to the unicellular Myzozoa, a clade that includes apicomplexan parasites and free-living dinoflagellates^22^. Attempts to delete the four genes using both genome-wide screens in *P. falciparum* and the murine malaria model parasite *Plasmodium berghei*, and targeted approaches in the latter, resulted in conflicting results^22–25^. Localization studies through fluorescent tagging of the endogenous *P. berghei* genes revealed a mitochondrial localization of three SPFH proteins throughout the *P. berghei* life cycle^22^. Although tagging of *PHB1* was unsuccessful, PHB1/2 heterodimerization is evolutionary well-conserved^15,26^ and *PfPHB1* ranks 84^th^ on the validated list of predicted *Plasmodium* mitochondrial proteins^27^ . Functional complementation of yeast *PHB* mutants provided further support for prohibitin heterodimerization in *P. falciparum*^28^. *Pf*PHBs were shown to be involved in stabilizing mitochondrial DNA, maintaining mitochondrial integrity, and rescuing yeast cell growth^28^. PHBL-deficient parasites failed to colonize *Anopheles* mosquitos as they arrest during ookinete development, which is correlated with depolarization of the mitochondrial membrane potential^22^.

The role and importance of STOML remains unclear. Genetic screens in *P. falciparum* and *P. berghei* both suggested a dispensable role, yet targeted approaches did never yield a pure isogenic or clonal line free of wild-type (WT) parasites, indicating possible developmental issues^22–24^ . Interestingly, *Pb*STOML localizes to punctate foci at the parasite mitochondrion during oocyst growth, often at organellar branching points.^22^ This specific mitochondrial localization combined with the uncertainty about its importance and function drove us to further investigate the role of STOML in the human malaria causing *P. falciparum*.

In this study, we show that deletion of *STOML* in *P. falciparum* causes a significant growth delay of asexual blood stages (ABS), while sexual-stage development is not affected. *Pf*STOML localizes to punctate foci at mitochondrial branch endings and at branching points throughout ABS development. *Pf*STOML resides in a large protein complex and pulldown experiments identified the metalloprotease FtsH as a likely interaction partner. We also show that the predicted AlphaFold Multimer structure of *Pf*STOML is highly similar to its bacterial family member HflK/C, which has recently been shown to form a large, oligomerized, vault structure around FtsH hexameres, this way regulating their accessibility. This suggests that a similar scenario might apply to the STOML-FtsH complex in *P. falciparum*. These results provide novel insights into the function of STOML in *P. falciparum* and pave the way for future studies into the function of SPFH proteins and their potential as antimalarial drug targets.

## Results

### Knockout of *Pf*STOML results in a significant growth defect

To study the function of *Pf*STOML (PF3D7_0318100) during ABS development, we aimed to generate *Pf*STOML knockout (KO) parasites using a targeted replacement strategy (Figure S1A). Although the first three transfection attempts were unsuccessful, we managed to generate two *Pf*STOML KO parasite lines in NF54 (*stoml*^-^) and the MitoRed background (*stoml*^_^_*mito*_), the latter harboring a fluorescent mitochondrial marker (mito-mScarlet)^29^. Correct integration and the absence of unaltered wild-type (WT) NF54 or MitoRed parasite contaminations were verified by diagnostic PCR (Figure S1B). To demonstrate if *PfSTOML* KO causes a growth defect, we set up a new competition growth assay analogous to the protocol used with *Plasmodium berghei*^30^. In both *Pf*STOML KO lines, *STOML* is replaced by *GFP* under the constitutive *P. falciparum* histone 2B (*Pf*H2B, PF3D7_1105100) promotor, making them green fluorescent. By mixing these with WT parasites harboring a constitutively expressed mScarlet, the relative abundance of red-only and green fluorescent parasites can be determined by flow cytometry and followed over time (Figure 1A). The average factor by which the red/green ratio changed from the first to the second timepoint in three independent experiments was defined as *f*_*r*_. We included a control condition in which mNeonGreen expressing WT parasites (*cyto-NG*) are mixed with mScarlet expressing WT parasites (*cyto-mScarlet*) (Figure S2). We confirmed that the ratio of red and green parasites in this control culture was stable over time (*f*_*r*_ = 1.0) (Figure 1B). However, when co-culturing either of our *PfSTOML* KO lines with cyto-mScarlet, we found that the red/green distributions shift significantly over time (fr = 20.9, p<0.0001). In one representative experiment, the ratio of *cyto-mScarlet* versus *stoml*^-^_*mito*_ parasites changes from 47:53 to 94:6 after one week and 99:1 after two weeks, indicating that *stoml*^-^_*mito*_ grows approximately 12 times slower in a week period compared to WT in mixed culture conditions. We observed a very similar trend in *stoml*^-^ mixed cultures. To test whether this growth defect is caused by lower number of viable offspring per parasite, reduced invasion, or delayed development throughout the ABS cycle, we quantified growth and analyzed stage development microscopically in two independent experiments, either every 8-16 h over an 88-h period or daily for eight days (Figure 1C-E). These experiments showed that *stoml*^-^_*mito*_ develops slower throughout the ABS replication cycle compared to MitoRed WT parasites. At the end of the first replication cycle (40 h), MitoRed WT cultures contained mostly segmented schizonts and rings, and the parasitemia has increased 6.0-fold compared to the 24 h timepoint (Figure 1D). *stoml*^-^_*mito*_ cultures contained mostly early schizonts and very few rings at 40 h, and the parasitemia has only increased 1.6-fold. However, at 48 h, *stoml*^-^_*mito*_ parasitemia has almost “caught up” with WT parasitemia with a 4.9-fold increase compared to the 24 h timepoint. This trend of delayed ABS development continues in the next replication cycles (Figure 1E). These results indicate that *stoml*^-^_*mito*_ has a growth defect that is mainly caused by slower and prolonged development throughout the asexual replication cycle.

**Figure 1.**
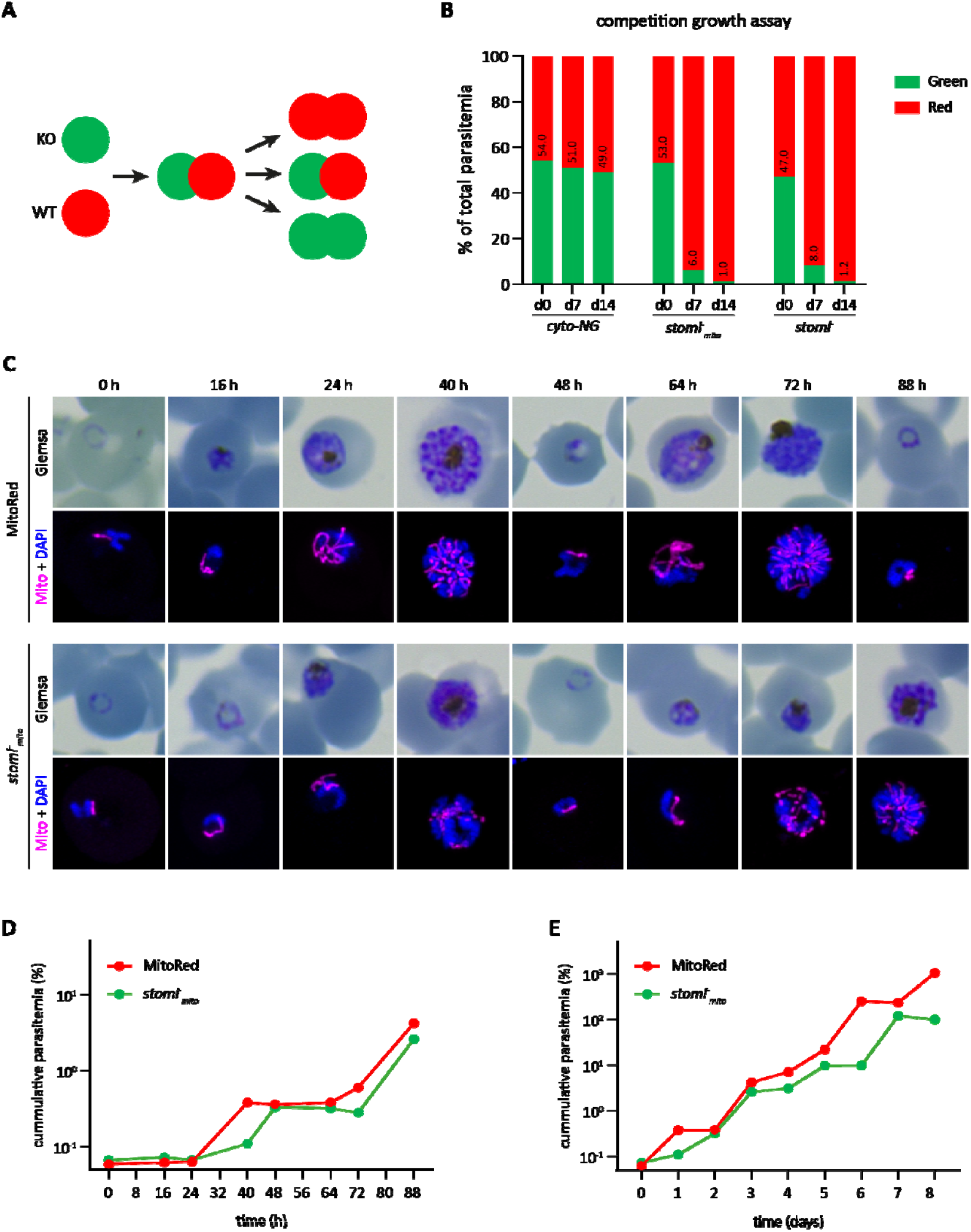
ABS development of stoml^_^_mito_ parasites is delayed. A) Schematic overview of the competition growth assay. PfSTOML knockout or control parasites expressing cytosolic GFP (green) are mixed with wild-type (WT) parasites expressing cytosolic mScarlet (red) in an approximate 1:1 ratio. Overtime, the distribution of red/green parasites is measured using flow cytometry. If PfSTOML knockout causes a growth defect, the WT population will grow faster and the ratio red/green parasites will shift. B) Bar graph showing distribution of red/green parasites in the competition growth assay at day 0, day 7, and day 14. The graph shows one representative example of three independent experiments. Cyto-mScarlet parasites are mixed with green cyto-NG (control), stoml, or stoml^-^ _mito_ parasites to create mixed cultures. C) Giemsa-stained thin blood smears and fluorescent images showing the mitochondrial mScarlet marker (magenta) and DNA (blue) in MitoRed (WT) and stoml _mito_ parasites over time. Fluorescence microcopy images are maximum intensity projections of Z-stack confocal Airyscan images. D-E) Growth curve of parasitemia of MitoRed and stoml^-^ _mito_ over time in hours (D) and days (E). To visualize continuous growth across culture dilutions, parasitemia values were corrected for dilution factors, resulting in cumulative (corrected) parasitemias exceeding 100%. The graphs show one representative experiments of two independent experiments.

-

### PfSTOML is unlikely to be involved in assembly of the respiratory chain

In other eukaryotes, stomatin-like proteins are thought to be involved in a variety of mitochondrial functions^13,20,31–33^. To explore if *Pf*STOML has a similar mitochondrial function in *P. falciparum*, we were curious to see if *Pf*STOML KO would alter mitochondrial dynamics. We compared mitochondrial morphology of *stoml*^-^_*mito*_ with MitoRed parasites and found no obvious differences throughout different stages of ABS development in two independent experiments (Figure S3). Mature *stoml*^*-*^*mito* schizonts showed divided and segregated mitochondria, similarly to MitoRed WT parasites.

SLP2, the human STOML homolog, has been indicated to play an essential role in the assembly of the respiratory chain^12,31^. To test if STOML has a similar function in *P. falciparum*, we investigated if *stoml*^-^ parasites would have an increased sensitivity to drugs targeting the respiratory chain as demonstrated successfully in the past for other mitochondrial proteins ^34-36^. We performed drug assays with different mitochondrial drugs (including DSM1, DSM265, atovaquone, ELQ300, and proguanil) and non-mitochondrial drugs (chloroquine, DHA, and MMV183) and found no difference in drug sensitivity between stoml and WT parasites in two independent experiments (Figure S4). Energy metabolism in *P. falciparum* ABS relies heavily on glycolysis and oxidative phosphorylation (OXPHOS) is only essential for ubiquinone recycling for pyrimidine synthesis^37^. However, in gametocytes, there is an increased TCA cycle utilization and presumably respiration^38,39^. To our surprise, *stoml*^-^_*mito*_ parasites develop to healthy-looking, mature gametocytes within a comparable time frame to WT parasites in three independent experiments. Furthermore, we found no obvious aberrations in mitochondrial morphology of mature male and female *stoml*^-^_*mito*_ gametocytes (Figure S5A). *stoml*^-^_*mito*_ parasites were still able to exflagellate and had dispersed mitochondria after activation, similarly to what we described for the MitoRed WT line^29^ (Figure S5B). Based on these results, it is unlikely that *STOML* is directly involved in maintaining mitochondrial morphology or respiratory chain assembly in *P. falciparum*.

### PfSTOML localizes to specific foci at the mitochondrion during ABS development

To learn more about the function of *Pf*STOML, we analyzed its subcellular localization. To do this, we generated a transgenic parasite line, *stoml-NG*, in which *STOML* is fused with a 3HA-mNG-GlmS tag. We also integrated a mitochondrial marker cassette mito-mScarlet for protein co-localization (Figure S1A). Correct integration and the absence of WT parasite contaminations were verified by diagnostic PCR (Figure S1B). Western blot analysis of ABS parasite extract confirmed expression of the full length *Pf*STOML-3HA-NG (Figure S1C). Live fluorescent microscopy of *stoml-NG* in three independent experiments showed localization of *Pf*STOML-3HA-NG to punctate foci during ABS development (Figure 2). In ring stages, *Pf*STOML-3HA-NG localized to a single spot, close to the mitochondrion (Figure 2A). As the parasites progress to late rings and the mitochondrion elongates, *Pf*STOML-3HA-NG is consistently found in two foci that reside at both endings of the mitochondrion (Figure 2A, 2B, Movie 1). In trophozoites, the mitochondrion starts to form a branched structure and the number of *Pf*STOML-3HA-NG foci per parasite increases (Figure 2C). *Pf*STOML-3HA-NG foci are found both at endings of mitochondrial branches, as well as branching points (Figure 2A). A similar localization is found in early schizont stages, where the mitochondrion forms a complex, branched network (Figure 2A, 2B, Movie 2). The number of foci per parasite increases during schizont maturation. In late schizonts, when the mitochondrial branches orient in a radial fashion prior to division, the *Pf*STOML-3HA-NG foci are found at the endings of most mitochondrial branches, although PfSTOML-3HA-NG signal can also be observed along mitochondrial branches (Figure 2A, 2B, Movie 3). In a fully segmented parasite that seems to have egressed form the RBC, only few *Pf*STOML-3HA-NG foci were found. The *Pf*STOML-3HA-NG foci are largely but not completely overlapping with the mito-mScarlet signal in trophozoite and schizont stages (Figure 2B), suggesting localization to the mitochondrial membrane, although we lack the resolution to distinguish between IMM and OMM. This unique localization pattern is different from the homogeneous mitochondrial localization of *Pb*STOML observed in *P. berghei* ABS^22^.

**Figure 2.**
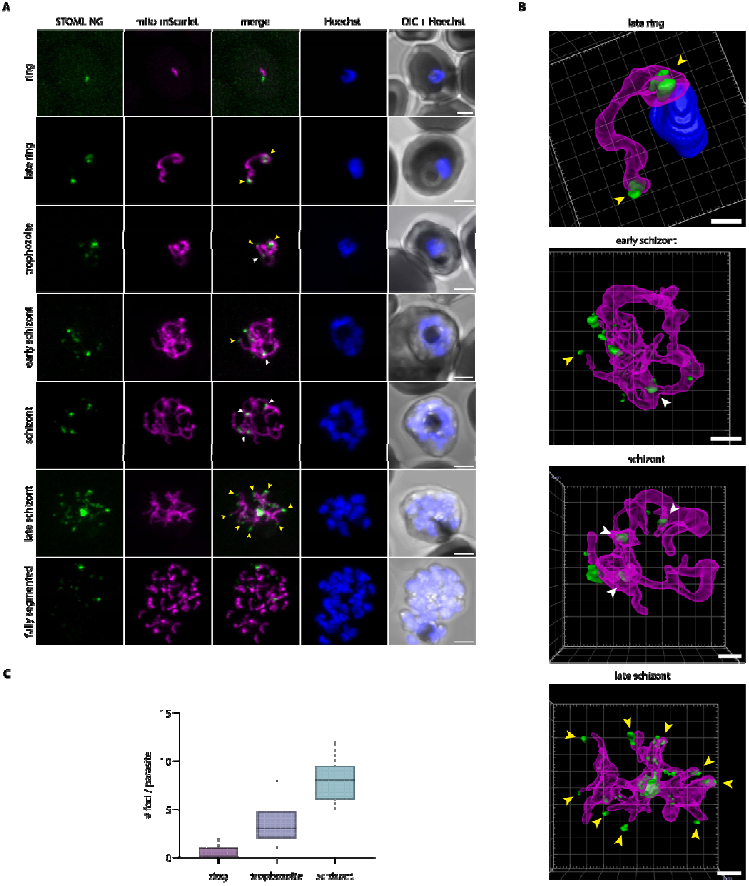
Localization of STOML-3HA-NG in ABS parasites. A) Live imaging of stoml-NG with PfSTOML-3HA-NG (green), mito-mScarlet mitochondrial marker (magenta), Hoechst for DNA visualization (blue), and DIC through the asexual replication cycle. Images are maximum intensity projections of Z-stack confocal Airyscan images. Arrowheads indicate PfSTOML-3HA-NG signal at mitochondrial branching points (white) or mitochondrial branch endings (yellow). Scale bars, 2 µm. B) 3D visualization of Z-stack confocal Airyscan images using Arivis 4D vision software. Fluorescent signal is segmented by manual thresholding. Arrowheads indicate STOML-NG signal at mitochondrial branching points (white) or mitochondrial branch endings (yellow). Scale bars, 1 µm. C) Boxplot indicating number of PfSTOML-3HA-NG foci per parasite in ring (n=7), trophozoite (n=12) and schizont (n=9) stages with a total of 28 parasites.

### *Pf*STOML resides in a large protein complex with *Pf*FtsH

PHBs and STOML are found in large hetero- and homo-oligomers at the IMM in other eukaryotes, such as humans and yeast^15,20^. *P. falciparum* mitochondrial complexome profiling data showed that PfSTOML migrates across a broad size range of ∼1.5-3.5 MDa on a native gel indicating that it forms part of one or potentially multiple large protein complexes (Figure 3A)^39^. In order to identify the proteins in complex with PfSTOML, we performed two independent pulldown experiments (Figure 3A-B, S7). In the first experiment, late-ABS *stoml-NG* parasites (24-40 h.p.i.) were lysed through saponin lysis and nitrogen cavitation, and the organelle fraction was used as input for co-immunoprecipitation with anti-mNG coated magnetic beads. As a control, the same fraction was loaded on uncoated beads. Mass spectrometry revealed that 27 proteins were significantly enriched after pulldown with mNG beads, of which PfSTOML was the most significantly enriched (Figure 3B, Table S2). For the second pulldown experiment, we generated a transgenic parasite line in which STOML is fused with an 3HA-GlmS tag, which we termed stoml-HA. Correct integration and the absence of WT parasite contaminations were verified by diagnostic PCR (Figure S1B) and western blot analysis confirmed expression of PfSTOML-3HA (Figure S1C). Fluorescence microscopy confirmed that PfSTOML-3HA localizes to mitochondrial branching points and branch endings, consistent with the pattern observed in live imaging of PfSTOML-3HA–mNG (Figure S1D, 2). This indicated that the addition of the large mNeonGreen tag does not affect the native mitochondrial localization of PfSTOML. The organelle fraction of late-ABS *stoml-HA* parasites was used as input for pulldown with anti-HA coated magnetic beads and empty protein G beads were used as control. In the second experiment, 122 proteins were significantly enriched after HA pulldown (Figure S7, Table S3). Three proteins were significant hits in both pulldown experiments: STOML, an ATP-dependent zinc metalloprotease FtsH (PF3D7_1464900), and a conserved protein of unknown function (PF3D7_1306200) (Figure 3C). In our published complexome profiling experiments, PF3D7_1306200 did not comigrate with STOML in either ABS or gametocyte stages (Figure 3A, S6)^39^ . PF3D7_1306200 is predicted to be an essential protein and is expressed in late schizonts^29,40,41^. It contains an AB-hydrolase domain and is thought to localize to the apicoplast ^42^. FtsH, on the other hand, is a predicted mitochondrial protein (ranking 265^th^)^27^ and phylogenetic analysis shows clustering with i-AAA proteases in the IMM^43^. Both in ABS and gametocytes, FtsH migrates corresponding to a mass of ∼2.5 MDa, which in gametocytes corresponds with the most dominant STOML migration peak (Figure 3A, S6)^39^ . In ABS, these normalized migration profiles show weaker apparent comigration, as FtsH overlaps with a minor peak of STOML. However, when considering absolute protein abundance (iBAQ values), the FtsH peak comigrates with a similar quantity of STOML, compatible with a putative complex (Figure S6). As STOML is substantially more abundant than FtsH (≈173-fold in gametocytes and ≈130-fold in ABS), this can obscure comigration in normalized profiles. The broad STOML migration pattern may therefore reflect multiple (sub)assemblies, with only a small subset of STOML associating with FtsH.

**Figure 3.**
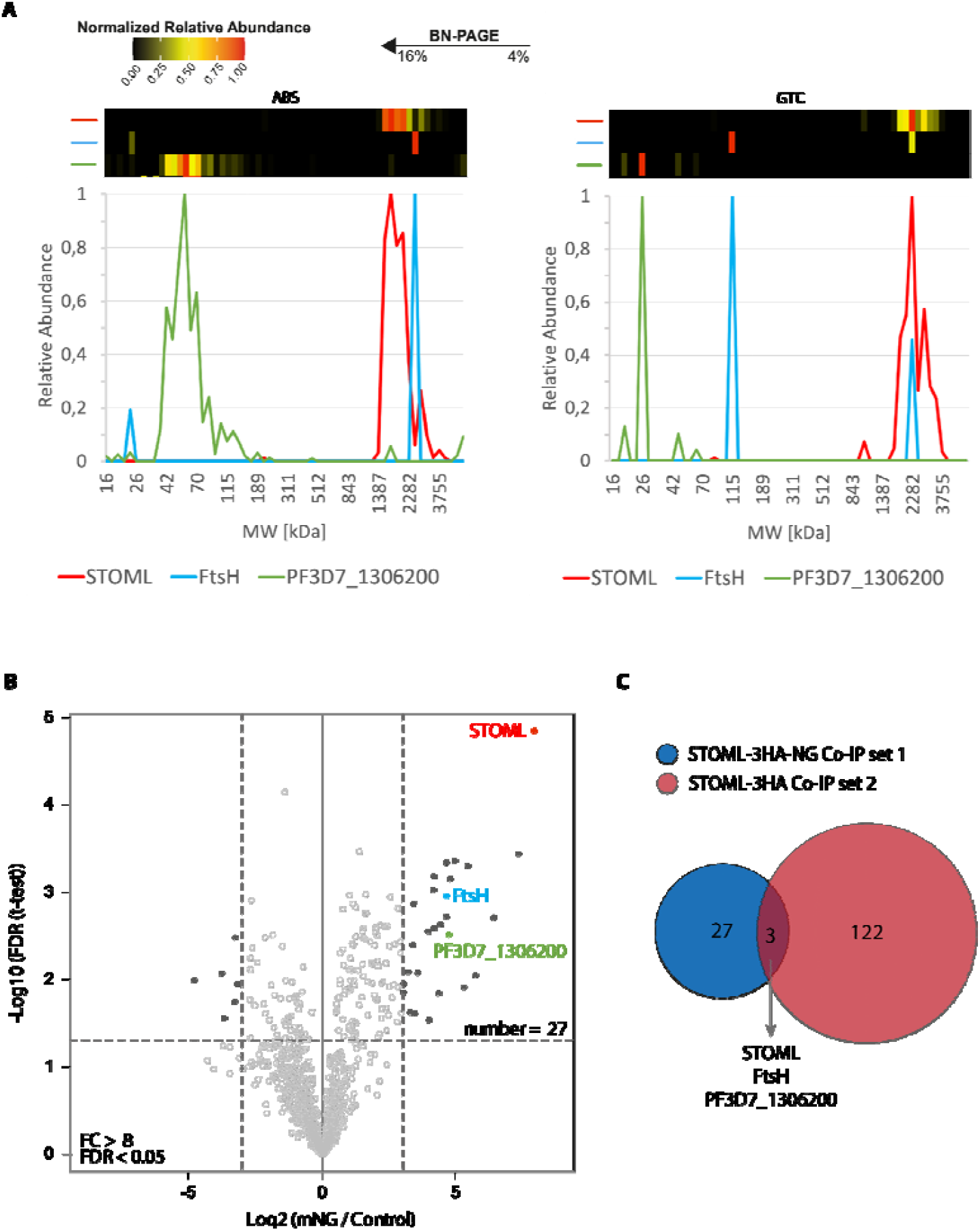
Identification and characterization of STOML protein complex. A) Heatmaps and line graphs based on previously published complexome profiling data^39^ showing migration patterns and relative protein abundance of STOML (red), FtsH (blue), and PF3D7_1306200 (green) in asexual blood stages (ABS) and gametocytes (GTC). Co-migration on blue native gel within the same molecular weight (MW) range (x-axis) indicates complex formation. Their relative abundance was normalized with the highest iBAQ value for a given protein set to 1 (shown red in the heatmap). B) Anti-mNG co-immunoprecipitation of PfSTOML-3HA-NG containing complexes. The volcano plot showing mean log_2_ fold changes (FC) and -log_10_ false discovery rate (FDR) for anti-HA pulldown in comparison with control pulldown. Horizontal and vertical dotted lines indicate log_2_ FC > 3 (FC > 8) and -log_10_ FDR > 1.301 (FDR > 0.05) respectively. Dark dots represent proteins that are highly enriched or reduced in the anti-mNG pulldown compared to the control pulldown. C) Overlap of enriched proteins in both anti-HA pulldown on STOML-HA and anti-NG pulldown on STOML-NG.

SPFH proteins are known to form large protein complexes with metalloproteases, regulating their protease activity^8,18,20,21,44^. The human STOML homolog, SLP2, forms a large proteolytic complex termed the SPY complex at the inner mitochondrial membrane with rhomboid protease PARL and i-AAA metalloprotease YME1L^20^. SLP2 regulates the activity of YME1L, which forms homo-hexamers and is involved in degradation of unfolded or excess mitochondrial proteins^45^. In *Trypanosoma brucei*, another unicellular protozoan parasite, SLP2 can also be found in a complex with the Yme1L homolog, *TbY*me1^46^. A BLAST search of hYME1L identified *P. falciparum* FtsH as top hit with 41% identity.

Cryo-electron microscopy revealed that the bacterial HflK and HflC form a large, hetero-oligomeric vault structure around four membrane-anchored FtsH hexamers (Figure 4E)^21^. We compared the predicted AlphaFold2^47^ structure of *Pf*STOML with the bacterial HflK/C complex (Figure 4A-D). Although *Pf*STOML is not predicted to contain a transmembrane domain^48^ the overall predicted structure of the protein is highly similar to its bacterial family members. To further explore if PfSTOML could form a similar multimer barrel structure, we used AlphaFold Multimer^49^ to predict the *Pf*STOML 24-multimer structure (Figure S8A). We used the SPFH2 and long alpha helix domains of *Pf*STOML, as these are the best predicted parts of the *Pf*STOML structure (pLDDT>90). The *Pf*STOML 24-multimer structure was predicted to form a distorted circular barrel structure with the endings of the multimer structure not joining together to close the structure, which visually seemed to include too many PfSTOML proteins (Figure S8A). In order to roughly estimate the correct amount of *Pf*STOML proteins in the barrel complex, we used AlphaFold Multimer to predict the structure of *Pf*STOML 8-multimer structure with the SPFH1, SPFH2, and long alpha helix domains (Figure S8B). We then measured the angle between the SPFH domains in the 8-multimer complex to estimate the curvature of the barrel (Figure S8C). We found an angle of 16.5 degrees between SPFH domains, suggesting that the *Pf*STOML complex might consist of approximately 22 *Pf*STOML monomers. AlphaFold Multimer predicts an intact, slightly oval barrel structure for a *Pf*STOML 22-multimer with SPFH2 and long alpha helix domains (Figure 4F, S8D). Considering the co-immunoprecipitation evidence and the high similarity between the predicted *Pf*STOML multimer structure and the HflK/C-FtsH complex, we hypothesize that STOML forms a similar supercomplex with FtsH likely in the IMM in *P. falciparum*.

**Figure 4.**
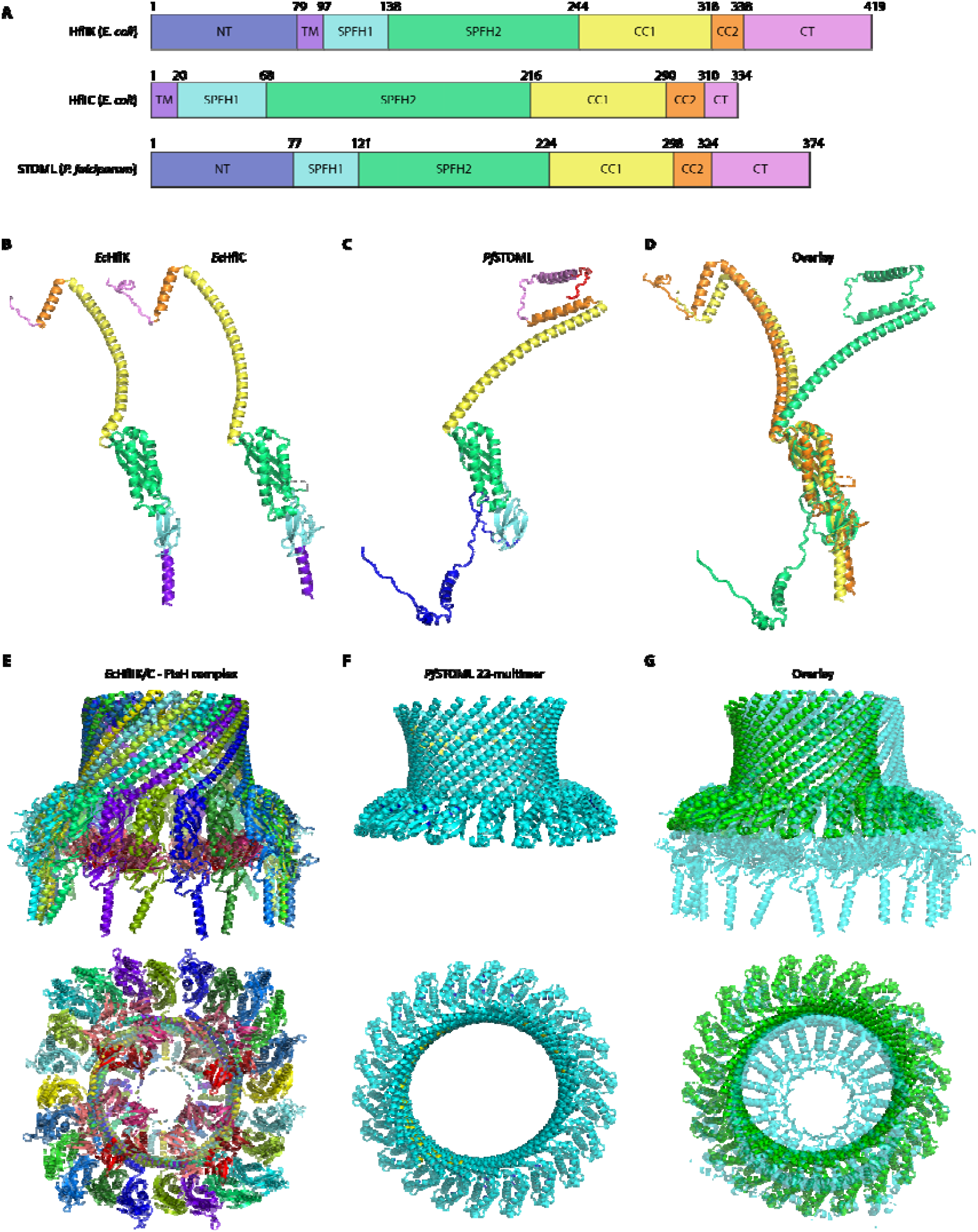
AlphaFold2 predictions of PfSTOML complex structure and comparison with the bacterial HflK/C supercomplex structures. A) Protein domains of EcHflK EcHflC, and PfSTOML, including the N-terminal domain (NT), transmembrane domain (TM), SPFH1 and SPHF2 domains, long coiled-coil domain 1 (CC1), coiled-coil domain 2 (CC2), and C-terminal region (CT). B) Protein structures of EcHflK and EcHflC determined by Ma et al.^21^ . C) Predicted AlphaFold2 structure of PfSTOML. D) Overlay of EcHflK and EcHflC structures with PfSTOML AlphaFold2 structure. E) Cryo-EM structure of HflK/C – FtsH complex with the side view (top) and bottom view (bottom). Purple/blue-colored proteins are HflC, green/yellow-colored proteins are HflK, red/pink-colored proteins are part of the FtsH hexamers^21^. PDB ID: 7VHP. F) Predicted AlphaFold Multimer structure of PfSTOML 22-multimer with side view (top) and top view (bottom), coloring according to model confidence with very high confidence (pLDDT > 90) dark blue, high confidence (pLDDT > 70) light blue, and low confidence (pLDDT > 50) yellow. G) Overlay of predicted AlphaFold Multimer PfSTOML 22-multimer structure (green) with HflK/C – FthsH complex (light blue) with side view (top) and top view (bottom).

## Discussion

The SPFH protein family is a highly conserved family involved in the formation of microdomains in membranes of various organelles. Prohibitins and stomatin-like proteins (STOML) are localized to the IMM and have been implicated in several mitochondrial functions, including cristae formation and assembly of the respiratory chain^11,12,14,31^. *Plasmodium* harbors four SPFH family members, including two prohibitins (PHB1 and PHB2), a prohibitin-like protein (PHBL), and STOML, which all localize to the mitochondrion^22^. STOML is likely essential in *P. berghei* and localizes to foci on mitochondrial branching points during oocyst stages. In this study, we investigated the function of STOML in the human malaria causing parasite *P. falciparum*.

In contrast to the *in vivo* murine model parasite *P. berghei* in which *STOML* could not be deleted, we were able to generate two *STOML* KO lines in *P. falciparum. stoml^-^* parasites developed slower throughout the ABS replication cycle compared to WT parasites, indicating an important but non-essential function for *Pf*STOML under *in vitro* culture conditions. In trophozoite and schizont stages, *Pf*STOML localizes to specific foci at the mitochondrial branching points and endings of mitochondrial branches. In late schizonts, when mitochondrial branches are oriented in a radial fashion prior to division^29^, PfSTOML has a punctate localization at the endings of mitochondrial branches. While STOML appears to have a more uniform mitochondrial distribution in most *P. berghei* life-cycle stages, *Pb*STOML localizes to foci on mitochondrial branching points in oocyst stages^22^. This specific localization suggests a potential function for *Pf*STOML in mitochondrial dynamics and/or segregation. On the other hand, *Pf*STOML knockout does not seem to affect mitochondrial division, and segregation (Figure S3). Therefore, the exact function of *Pf*STOML at this specific mitochondrial localization remains to be elucidated.

In human, yeast and plants, STOML has been indicated to play a role in the assembly of the respiratory chain^12,13,31^. We hypothesized that STOML might have a similar function in *P. falciparum* and that *Pf*STOML knockout would lead to an increased sensitivity to drugs targeting the respiratory chain, such as atovaquone and ELQ300. Such an approach has been used successfully in several studies of mitochondrial proteins in *P. falciparum* highlighting potential roles in respiratory chain assembly or functioning of *Pf*Rieske, *Pf*PNPLA2, and *Pf*OPA3^34–36^ . However, our data show no changes in drug sensitivity upon *Pf*STOML knockout. Additionally, *stoml*^-^_*mito*_ parasites were able to form healthy gametocytes and undergo male gametogenesis, during which mitochondrial ATP synthesis is thought to be essential^50^. This indicates that in contrast with orthologs in other eukaryotes, *Pf*STOML appears not directly involved in respiratory chain assembly in *P. falciparum*.

Our mitochondrial complexome profiling data show that PfSTOML consistently resides in one or more large ∼1.5-3.5 MDa protein complexes^39^. SPFH proteins are known to form large homo- or heteromeric complexes, often together with proteases^8,18^. In order to further characterize STOML complexes in *P. falciparum*, we performed two complementary STOML-pulldown experiments to identify potential interactors. Three proteins, including *Pf*STOML, *Pf*FtsH metalloprotease, and a protein of unknown function (PF3D7_1306200), were identified as significantly enriched in both pulldown experiments. PF3D7_1306200 contains an AB-hydrolase domain and is predicted to be essential^40^. Similarly to *Pf*STOML, it is mostly expressed in schizont stages^41^, yet the putative hydrolase is predicted to localize to the apicoplast^42,51^ and did not show comigration with *Pf*STOML on a native gel in our complexomics data^39^. Of note, for the co-immunoprecipitation experiments we used synchronized late-stages samples whereas the complexome profiling was performed on mixed ABS. Based on its predicted apicoplast localization, and the lack of comigration with *Pf*STOML on a native gel, it seems unlikely that PF3D7_1306200 forms a complex with *Pf*STOML at the IMM, yet, the consistent pull down with the *Pf*STOML is remarkable^52^. Reciprocal pull-down experiments could confirm interaction between PF3D7_1306200 and *Pf*STOML. During asexual and sexual blood-stage development there are plentiful of close appositions of the mitochondrion and apicoplast, the nature of which remains enigmatic ^29^. It might be possible that the *Pf*STOML - PF3D7_1306200 interaction plays a role in these organelle contact sites. A more detailed microscopic analysis of contact sites with (conditional) knockout of both proteins could shed more light on this hypothesis.

*Pf*FtsH belongs to the AAA (<u>A</u>TPases <u>A</u>ssociated with various cellular <u>A</u>ctivities) metalloprotease family at the IMM, which play a role in protein surveillance by degrading non-native integral membrane proteins and membrane associated proteins such as unassembled units of the respiratory chain^52,53^. *P. falciparum* harbors three FtsH homologs^43^. Two of the *P. falciparum* FtsH homologs, including PF3D7_1464900, which we identified in our *Pf*STOML pull-down, locate to the IMM, have a single transmembrane domain, and cluster with i-AAA FtsH homologs, which are exposed to the intermembrane space. The third homolog, *Pf*FtsH1 (PF3D7_1239700), has two transmembrane domains and clusters with *m-AAA* FtsH homologs at the IMM that are exposed to the mitochondrial matrix. *Pf*FtsH1 forms oligomeric complexes and has a punctate distribution on the mitochondrial branching points in late trophozoite and early schizont stages^43^, which is similar to *Pf*STOML distribution in these stages. Contradictory, another study showed that actinonin, a small molecule inhibitor, targets *Pf*FtsH1 and disrupts apicoplast biogenesis^54^ and its homolog in *T. gondii* is also localized to the apicoplast^55^. Expression of *Pf*FtsH1 in *E. coli* causes defective cytokinesis, implying a potential role in organelle division. Unfortunately, Amberg-Johnsen and colleagues were unable to generate endogenously tagged knockdown parasite lines of the two *i-AAA* proteases^54^. Therefore, the exact function and substrates of FtsH in *P. falciparum* remain to be elucidated.

The interaction between *Pf*STOML and *Pf*FtsH is well-supported by evidence. Their respective human homologs, SLP2 and Yme1L, form the SPY complex^20^, which is essential for the proteolytic regulation of proteins involved in mitochondrial dynamics and quality control. Yme1L also contributes to OPA1 cleavage, a mitochondrial GTPase which is involved in mitochondrial fusion and cristae formation^56,57^, however, no *Plasmodium* OPA1 homolog has been identified to date. In *Trypanosoma brucei*, TbSLP2 forms a large mitochondrial complex with TbYme1, which is involved in mitochondrial stress resistance^46^. Furthermore, in the filamentous fungus *Neurospora crassa*, STOML2 has been found in a large complex with an *i*-AAA protease (IAP1)^58^. Other SPFH family members are also known to form large complexes with AAA+ proteases, such prohibitins with Yta10/Yta12 in yeast^8^ or HflK/C with FtsH in bacteria^18^. Characterization of the HflK/C-FtsH supercomplex structure showed that

HflK/C forms a 24-heteromer vault structure around four hexameric FtsH complexes at the bacterial membrane^21^ (Figure 4E). Here, we show that the *Pf*STOML AlphaFold2 predicted structure shows high similarity with that of bacterial HflK/C. Our AlphaFold Multimer predictions suggest that *Pf*STOML might form a 22-multimer barrel structure that is highly similar to the vault structure of HflK/C-FtsH supercomplex in bacteria. Although these predictions are based on modelling and therefore need to be interpreted with caution, the high structural similarity of the *Pf*STOML 22-multimer complex with HflK/C-FtsH supercomplex, combined with our co-immunoprecipitation data, suggests that *Pf*STOML might form a similar supercomplex structure with *Pf*FtsH, possibly regulating *Pf*FtsH accessibility.

Taken together, knockout of *Pf*STOML causes a significantly delayed ABS development, while gametocytes develop normally. *Pf*STOML has a punctate distribution to mitochondrial branching points and endings of mitochondrial branches but knockout of *Pf*STOML does not affect mitochondrial morphology. Knockout of *Pf*STOML did not affect sensitivity to drugs targeting the respiratory chain, suggesting that *Pf*STOML is not directly involved in respiratory chain assembly. *Pf*STOML resides in a large supercomplex with PfFtsH, likely forming a large, multimeric barrel structure that regulates the accessibility of *Pf*FtsH, similar to its bacterial family members. Although the exact function of the STOML-FtsH complex in *P. falciparum* remains to be elucidated, these results could pave the way for future studies into this highly conserved protein family and their role in proteolytic processes and membrane organization.

## Materials and Methods

### *P. falciparum* culture and transfections

*P. falciparum* NF54 and mutant parasites lines were cultured in RPMI1640 medium supplemented with 25 mM HEPES, 10% human type A serum (Sanquin, The Netherlands) and 25 mM NaHCO_3_ (complete medium). Parasites were cultured in 5% human RBCs type O (Sanquin, The Netherlands) at 37°C with 3% O_2_ and 4% CO_2_. For transfection, 60 μg of HDR plasmid was linearized by overnight digestion, precipitated, and transfected with 60 μg Cas9 plasmid using either RBC loading or ring transfection^59^ . For RBC loading, plasmids were loaded into RBCs by electroporation (310 V, 950 μF) and a trophozoite parasite culture was added to the transfected RBCs. One day after transfection, parasites were treated with 2.5 nM WR99210 (Jacobus Pharmaceutical) for five days. For ring transfection, a ring-stage sorbitol synchronized parasite culture was transfected with the plasmids by electroporation (310 V, 950 μF). Five hours after transfection, parasites were treated with 2.5 nM WR99210 for five days. Success of transfection was assessed by diagnostic PCR (Figure S1). Gametocyte cultures were maintained in a semi-automatic culturing system with media changes twice a day^60^ . Gametocytes were stress-induced through asexual overgrowing. A mixed asexual culture of 1% was set up and cultured for up to 2 weeks.

### Plasmid constructs

To generate the *STOML* KO repair plasmid, the pGK plasmid was used, which contains a pBAT backbone^61^ with H2B promotor, GFP and PBANKA_142660 bidirectional 3’UTR, flanked by multiple cloning sites. The 5’ and 3’ homology regions (HRs) were amplified and cloned into pGK sequentially, using XmaI + XhoI and NcoI + EcoRI restriction sites, respectively, generating pRF0038 *STOML* KO repair plasmid. CRISPR-Cas9 guide plasmids targeting two different sites in STOML were generated. Guide oligonucleotides were annealed and cloned into pMLB626 plasmid^62^ (a kind gift from Marcus Lee) using BbsI restriction enzyme, generating the pRF0039 and pRF0040 final guide plasmids (Table S1).

To generate *STOML* tagging repair plasmid, pRF0079 empty tagging plasmid was used, containing 3HA-NG-GlmS, PBANKA_142660 bidirectional 3’UTR, and the mito-mScarlet mitochondrial marker^29^ . 5’ HR was generated by overlap PCR, harboring a shield mutation that prevents cutting of CRISPR-Cas9 when the construct is integrated. 5’ and 3’ HRs were cloned into pRF0079 using KpnI + BamHI and EcoRI + NgoMIV restriction enzymes, respectively, generating pRF0166 *STOML* tagging plasmid. Because this plasmid was unsuccessful in generating mutant parasite line after three transfection attempts, we decided to clone the *DHFR* selection marker in the repair plasmid and remove it from the guide plasmid. By addition of WR after transfection, we will then directly select for parasites with integration of the *DHFR* cassette, instead of selection on the guide plasmid. The *DHFR* cassette was removed from pRF0040 guide plasmid by digestion with EcoRI and ApaI, followed by blunt end generation with DNA polymerase I (klenow), following manufacturer’s instructions, and ligation. The new guide plasmid without *DHFR* was termed pRF0210. The *DHFR* cassette cloned from MLB626 plasmid into pRF0166 with SphI and EcoRI restriction enzymes, generating pRF0213 3HA-NG-GlmS tagging plasmid with mito-mScarlet and *DHFR* selection marker. Since a big fluorescent tag might interfere with protein function, we also generated a *STOML* tagging repair plasmid by removing mNG from pRF0213, generating pRF0266 3HA-GlmS tagging plasmid, using BamHI and NheI restriction enzymes.

For generation of the repair plasmids for cyto-mScarlet and cyto-mNG parasite lines (used for the competition growth assay), SIL7 reporter plasmid (pRF0057) was used^29^ . *mScarlet* was amplified from p1.2RhopH3-HA-mScarlet^63^ (a kind gift from Prof. Alan Cowman) (Table S1) and cloned into pRF0057 using AfeI and NheI restriction enzymes, generating pRF0278 cyto-mScarlet repair plasmid. *mNeonGreen* was amplified from pRF0079 plasmid and cloned into pRF0278 with AflII and NheI restriction sites to generate pRF0290, the cyto-mNG repair plasmid. CRISPR-Cas9 guide plasmids targeting SIL7 were used^29^ . All enzymes used were obtained via New England Biolabs.

### Competition growth assay

For the competition growth assay, parasite lines harboring a cytosolic mScarlet or cytosolic mNG were generated by integration of cyto-mScarlet and cyto-mNG constructs in SIL7 integration site^29^ . Cyto-mScarlet, Cyto-mNG, *stoml*^-^ and *stoml*^-^_*mito*_ were synchronized by a 63% Percoll centrifugation. Late-stage parasites were isolated from the Percoll gradient and added to fresh RBCs. Four hours later, 5% sorbitol synchronization was performed, which allowed only young rings that just invaded a new RBC to survive. Ring-stage parasites were counted and diluted to each have 0.4% final parasitemia in Cyto-mScarlet + Cyto-mNG, Cyto-mScarlet + *stoml*^-^, and Cyto-mScarlet + *stoml*^-^_*mito*_ mixed cultures. Samples for flow cytometry analysis were taken directly after set-up, at day 7 and at day 14. Samples from each mixed culture were taken and stained with 0.5 μg/ml Hoechst 33342 (Invitrogen, H3570) for 30 min at 37°C. Samples were directly analyzed on BD FACSAria™ III Cell Sorter and number of red and green parasites were counted. Data was analyzed in FlowJo (version 10.10).

### Growth assay

For this growth assay, MitoRed (WT), *stoml*^-^, and *stoml*^-^_*mito*_ parasites were synchronized by a 63% Percoll centrifugation. Late-stage parasites were isolated from the Percoll gradient and added to fresh RBCs. Four hours later, 5% sorbitol synchronization was performed, which allowed only young rings that just invaded a new RBC to survive. Ring-stage parasites were counted and diluted to 0.05% parasitemia. Samples for flow cytometry analysis and fluorescent microscopy were taken directly after setup (t=0), and then every 8, 16 or 24 h over a period of 8 days. To prevent overgrowth, parasite cultures were cut back 1/100 on day 3, and 1/50 on day 6. For flow cytometry, samples were taken and fixed in 0.25% glutaraldehyde. All samples from different time points were processed at the same time, by staining with Hoechst 33342 for 30 min at 37°C and then analyzed by flow cytometry (Beckman Coulter Cytoflex) to determine parasitemia using the 405 nm laser. Data was analyzed in FlowJo (version 10.10). Final parasitemia was adjusted for the dilution factor, explaining why final parasitemia can reach more than 100%. For fluorescent microscopy, parasite samples were processed as described in “fixed imaging” paragraph below.

### Live imaging

*Stoml-NG* parasites were stained with Hoechst 33342 for 30 min at 37°C and settled in an 8-well imaging chamber (Ibidi) in complete media without phenol red. Parasites were imaged on a Zeiss LSM880 or LSM900 Airyscan microscope with 63x oil objective and 37°C heated stage, using 405, 488, and 561 nm excitation lasers. Images were Airyscan processed before analysis with FIJI software^64^ and Arivis vision4D (Zeiss).

### Fixed imaging

For fixed immunofluorescence microscopy of the asexual and sexual blood stages of the *stoml*^-^_*mito*_ lines, parasites were settled on a poly-L-lysine coated coverslip for 20 min at 37°C. Parasites were fixed using 4% EM-grade paraformaldehyde and 0.0075% EM-grade glutaraldehyde in PBS for 20 min at room temperature. Cells were permeabilized with 0.1% Triton X-100 for 10 min, before staining using 1 μM DAPI in PBS for 1 h. For imaging of *stoml*^-^_*mito*_ gametocytes, parasites were stained for 1 h with primary mouse anti-alfa-tubulin antibody (A11126, ThermoFisher, 1:500) and secondary goat anti-mouse Alexa Fluor 647 (A21247, ThermoFisher, 1:200) antibody in 3% BSA in PBS. For experiments using the stoml-HA parasite line, asexual blood stage cultures were stained with MitoBrilliant™ 646 (Tocris, 1:10,000 in complete medium) for 20 min at 37°C. Afterwards, cells were fixed and permeabilized as described above. Parasites were stained for 1 h with primary rat anti-HA (3F10, Roche, 1:100) and secondary goat anti-rat Alexa Fluor 488 (A48262, ThermoFisher, 1:200) antibodies diluted in 3% BSA in PBS, followed by a DAPI staining as described above. All slides were mounted with Vectashield (Vector Laboratories). Samples were imaged with a Zeiss LSM880 or LSM900 Airyscan microscope with 63x oil objective and 405, 488, 561, and 633 nm excitation lasers. Images were Airyscan processed using Zeiss Zen Blue Software, before analysis with FIJI software^64^ and Arivis vision4D (Zeiss).

### Co-immunoprecipitation assay

*Stoml-HA* and *stoml-NG* parasites were synchronized with 5% sorbitol and harvested 22 h later to obtain late-stage parasites. Parasites were treated with 0.06% saponin, snap-frozen in liquid nitrogen, and C until further processed. Nitrogen cavitation was used for cell disruption as described stored at -80° 39. On the day of the experiment, 18 pellets of 30-ml cultures per parasite line were resuspended and pooled in 25 ml ice-cold MESH-buffer (250 mM sucrose, 10 mM HEPES, 1 mM EDTA, 1× cOmplete™ EDTA-free Protease Inhibitor Cocktail (Sigma), pH 7.4). The sample was added to the pre-chilled cell disruption vessel (#4639 Parr Instrument Company) and pressurized with nitrogen gas at 1500 psi for 45 min on ice. The parasites were then sheared through slow release. The organelle-enriched fraction was obtained by differential centrifugation as described^39^ . Protein concentrations were determined by Pierce™ BCA Protein Assay Kit (Thermo Scientific). Samples were solubilized with n-dodecyl-β-D-maltoside (DDM) (Sigma), using 3:1 detergent:protein (w/w) ratios. Solubilized samples were spun down at 22,000 x g at 4°C. Supernatant derived from *stoml-NG* samples were applied on ChromoTek mNeonGreen-Trap magnetic agarose beads (ChromoTek), or empty binding control agarose beads (ChromoTek).C Supernatant from stoml-HA samples were applied on PierceTM HA-tag magnetic beads (Thermofisher) or empty protein G binding control beads (Thermofisher). Both pulldowns were carried out with three technical replicates. Beads were incubated at 4°C for 30 minutes with gentle agitation and then washed twice with washing buffer (PBS, 1mM EDTA, 1× cOmplete™ EDTA-free Protease Inhibitor Cocktail, 0.05% DDM) and three times with ice-cold PBS, using a magnetic stand. After washes, on bead digestion was performed as follows: beads were resuspended in 50 μl elution buffer (2M urea, 100 mM Tris-HCl pH 8.0, 10 mM DTT) and incubated for 20 minutes at 25°C while shaking. To alkylate cysteines, iodoacetamide was added to a final concentration of 50 mM. Samples were kept in the dark for 10 min at 25°C. Subsequently, 0.25 μg of sequencing grade tryspin (Promega) was added to digest the proteins. The samples were shaken at 25°C for 2 h. The supernatants, containing the partially digested proteins, were collected and 50 μl of fresh elution buffer was added to the beads and shaken for another 5 min. Next, these supernatants were collected and combined with the first supernatant. Another 0.1 μg of trypsin was added, to stimulate overnight digestion at 25°C. The next day, samples were concentrated and purified on C18 stagetips ^65^ . Samples were analyzed on a Thermo Exploris 480 mass spectrometer, operated with an online Easy-nLC 1000. A gradient of buffer B (80% acetonitrile, 0.1% formic acid) was applied for 60 min. The mass spectrometer was ran in Top20 mode, while dynamic exclusion was enabled for 45 sec. Raw data was analyzed using Maxquant version 1.6.6.0^66^ with a Plasmodium database (strain 3D7, version August 5th 2022, obtained from plasmodb.org^67^). LFQ, iBAQ and match between runs were enabled, and deamidation (NQ) was added as additional variable modification. The output was filtered using Perseus 1.5.0.15^68^ . Proteins marked as potential contaminants, reverse hits, and proteins with less than 2 peptides were removed. Samples were grouped into triplicates, and proteins with less than 3 valid values in at least 1 group were removed, after which missing values were imputed using the default settings. A t-test was performed to identify specific outliers. Data was visualized using R. The mass spectrometry proteomics data have been deposited to the ProteomeXchange Consortium via the PRIDE^69^ partner repository with the dataset identifier PXD039772.

### Drug sensitivity assay

NF54 and *stoml*^-^ parasites were used in a replication assay as described^70^ to determine sensitivity to anti-malarial compounds. Briefly, parasites were diluted to 0.83% parasitemia and 3% hematocrit. Thirty microliters of diluted parasites were combined with 30 µl of compound serially diluted in dimethyl sulfoxide (DMSO) and RPMI 1640 medium to reach a final DMSO concentration of 0.1% in a total assay volume of 60 µl. Parasites were incubated at 37°C for 72 hours with mixed gas (3% O_2_ and 4% CO_2_). Then 30 µl of lysis buffer containing 1:15,000 SYBR Green reagent (Life Technologies), 13.3 mM Tris-HCl, 3.3 mM EDTA, 0.067% TritonX-100 and 0.0053% saponin was added and fluorescence intensity was quantified using BioTek Synergy 2 plate reader. GraphPad Prism was used for data analysis and inhibitory dose-response curves were determined with a variable slope model, in which the curve is generated with the following formula: 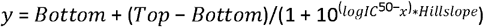.

### DALI search and AlphaFold2 structure predictions

AlphaFold Multimer49 predictions were performed using the COSMIC^2^ platform (https://cosmic-cryoem.org/tools/alphafoldmultimer/). All predicted protein and protein complex structures and alignments were visualized using PyMOL Molecular Graphics System (version 2.5.2. Schrödinger, LLC).

## Supporting information

Response to reviewers

## Acknowledgements

We are very grateful to Akhil Vaidya for the insightful discussions. We would like to thank all members of the molecular & cellular parasitology team at Radboudumc for the insightful discussions. We would like to thank the Radboud Technology Center Microscopy for use of their microscopy facilities. We are grateful to Michiel Vermeulen and his team for providing the mass spectrometry facility and to Dick Zijlmans for proofreading the manuscript. We also thank the Radboud Technology Center Flowcytometry for the use of their flowcytometry facilities. We are grateful to Koen Dechering and Tonnie Huijs for their support with the drug sensitivity assays. We thank Marcus Lee for providing the pMLB626 CRISPR/Cas9 guide plasmid and Richárd Bártfai for suggesting using the H2B promoter.

## Financial disclosure

This work was supported by the Netherlands Organisation for Scientific Research (NWO-VIDI864.13.009 to FE and TWAK) and the Radboud University Medical Center (Radboudumc Individual PhD for Masters round 2018 to JMJV; Radboudumc Individual PhD for Masters round 2021 to EB; RIMLS018-009b to CB and TWAK). The funders had no role in study design, data collection and analysis, decision to publish, or preparation of the manuscript.

## Author Contributions

JV and TK designed the study. JV, CB, MH, and NP generated transgenic parasite lines. JV and EB conceived and performed experiments. CB and MH developed the competition growth assay. FE reanalysed the complexome data. CS performed the mass spectrometry experiments and data analysis. JV wrote the first draft of the manuscript. EB and TK edited the manuscript. TK supervised the project. All authors read and approved of the manuscript.

## Notes

### Competing Interest Statement

The authors have declared no competing interest.

### Summary of Updates

Corrections to author names and contributions and addition of a financial disclosure.

## References

1. WHO. World Malaria Report 2023. (2023).

2. Lamb, I. M., Okoye, I. C., Mather, M. W. & Vaidya, A. B. Unique Properties of Apicomplexan Mitochondria. Annu Rev Microbiol 77, 541–560 (2023).

3. Goodman, C. D., Buchanan, H. D. & McFadden, G. I. Is the mitochondrion a good malaria drug target? Trends Parasitol 33, 185–193 (2017).

4. Laude, A. J. & Prior, I. A. Plasma membrane microdomains: Organization, function and trafficking (Review). Mol Membr Biol 21, 193–205 (2004).

5. Whitelegge, J. Up close with membrane lipid-protein complexes. Science (1979) 334, 320–321 (2011).

6. Browman, D. T., Hoegg, M. B. & Robbins, S. M. The SPFH domain-containing proteins: more than lipid raft markers. Trends Cell Biol 17, 394–402 (2007).

7. Tavernarakis, N., Driscoll, M. & Kyrpides, N. C. The SPFH domain: Implicated in regulating targeted protein turnover in stomatins and other membrane-associated proteins. Trends Biochem Sci 24, 425–427 (1999).

8. Steglich, G., Neupert, W. & Langer, T. Prohibitins regulate membrane protein degradation by the m-AAA protease in mitochondria. Mol Cell Biol 19, 3435–3442 (1999).

9. Merkwirth, C. & Langer, T. Prohibitin function within mitochondria: Essential roles for cell proliferation and cristae morphogenesis. Biochim Biophys Acta Mol Cell Res 1793, 27–32 (2009).

10. Oyang, L. et al. The function of prohibitins in mitochondria and the clinical potentials. Cancer Cell Int 22, 1–10 (2022).

11. Merkwirth, C. et al. Prohibitins control cell proliferation and apoptosis by regulating OPA1-dependent cristae morphogenesis in mitochondria. Genes Dev 22, 476–488 (2008).

12. Da Cruz, S. et al. SLP-2 interacts with prohibitins in the mitochondrial inner membrane and contributes to their stability. Biochim Biophys Acta Mol Cell Res 1783, 904–911 (2008).

13. Christie, D. A. et al. Stomatin-Like Protein 2 Binds Cardiolipin and Regulates Mitochondrial Biogenesis and Function. Mol Cell Biol 31, 3845–3856 (2011).

14. Mitsopoulos, P. et al. Stomatin-Like Protein 2 Is Required for In Vivo Mitochondrial Respiratory Chain Supercomplex Formation and Optimal Cell Function . Mol Cell Biol 35, 1838–1847 (2015).

15. Nijtmans, L. G. J. et al. Prohibitins act as a membrane-bound chaperone for the stabilization of mitochondrial proteins. EMBO Journal 19, 2444–2451 (2000).

16. He, J. et al. Mitochondrial nucleoid interacting proteins support mitochondrial protein synthesis. Nucleic Acids Res 40, 6109–6121 (2012).

17. Mitsopoulos, P. et al. Stomatin-like protein 2 deficiency results in impaired mitochondrial translation. PLoS One 12, 1–13 (2017).

18. Kihara, A., Akiyama, Y. & Ito, K. A protease complex in the Escherichia coli plasma membrane: HflKC (HflA) forms a complex with FtsH (HflB), regulating its proteolytic activity against SecY. EMBO Journal 15, 6122–6131 (1996).

19. Osman, C., Merkwirth, C. & Langer, T. Prohibitins and the functional compartmentalization of mitochondrial membranes. J Cell Sci 122, 3823–3830 (2009).

20. Wai, T. et al. The membrane scaffold SLP2 anchors a proteolytic hub in mitochondria containing PARL and the i □ AAA protease YME1L . EMBO Rep 17, 1844–1856 (2016).

21. Ma, C. et al. Structural insights into the membrane microdomain organization by SPFH family proteins. Cell Res 32, 176–189 (2022).

22. Matz, J. M., Goosmann, C., Matuschewski, K. & Kooij, T. W. A. An Unusual Prohibitin Regulates Malaria Parasite Mitochondrial Membrane Potential. Cell Rep 23, 756–767 (2018).

23. Zhang, M. et al. Uncovering the essential genes of the human malaria parasite Plasmodium falciparum by saturation mutagenesis. Science 360, eaap7847 (2018).

24. Bushell, E. et al. Functional Profiling of a Plasmodium Genome Reveals an Abundance of Essential Genes. Cell 170, 260–272.e8 (2017).

25. Jain, S., Narwal, M., Omair Anwar, M., Prakash, N. & Mohmmed, A. Unravelling the anti-apoptotic role of Plasmodium falciparum Prohibitin-2 (PfPhb2) in maintaining mitochondrial homeostasis. Mitochondrion 79, (2024).

26. Artal-Sanz, M. & Tavernarakis, N. Prohibitin and mitochondrial biology. Trends Endocrinol Metab 20, 394–401 (2009).

27. Esveld, S. L. Van et al. A Prioritized and Validated Resource of Mitochondrial Proteins in Plasmodium Identifies Unique Biology. mSphere 6, e00614–21 (2021).

28. Saini, M. et al. Characterization of Plasmodium falciparum prohibitins as novel targets to block infection in humans by impairing the growth and transmission of the parasite. Biochem Pharmacol 212, 115567 (2023).

29. Verhoef, J. M. J. et al. Detailing organelle division and segregation in Plasmodium falciparum. Journal of Cell Biology 223, (2024).

30. Matz, J. M. et al. The Plasmodium berghei translocon of exported proteins reveals spatiotemporal dynamics of tubular extensions. Sci Rep 5, 1–14 (2015).

31. Gehl, B. & Sweetlove, L. J. Mitochondrial Band-7 family proteins: Scaffolds for respiratory chain assembly? Front Plant Sci 5, 1–6 (2014).

32. Heredia, M. Y. & Rauceo, J. M. The spfh protein superfamily in fungi: Impact on mitochondrial function and implications in virulence. Microorganisms 9, (2021).

33. Tondera, D. et al. SlP-2 is required for stress-induced mitochondrial hyperfusion. EMBO Journal 28, 1589–1600 (2009).

34. Sheokand, P. K., Pradhan, S., Maclean, A. E., Mühleip, A. & Sheiner, L. Plasmodium falciparum Mitochondrial Complex III, the Target of Atovaquone, Is Essential for Progression to the Transmissible Sexual Stages. Int J Mol Sci 25, (2024).

35. Pietsch, E. et al. A patatin-like phospholipase is important for mitochondrial function in malaria parasites. mBio 14, (2023).

36. Narwal, M., Jain, S., Rathore, S. & Mohmmed, A. Plasmodium falciparum OPA3-like protein (PfOPA3) is essential for maintenance of mitochondrial homeostasis and parasite proliferation. FASEB Journal 37, 1–14 (2023).

37. Painter, H. J., Morrisey, J. M., Mather, M. W. & Vaidya, A. B. Specific role of mitochondrial electron transport in blood-stage Plasmodium falciparum. Nature 446, 88–91 (2007).

38. MacRae, J. I. et al. Mitochondrial metabolism of sexual and asexual blood stages of the malaria parasite Plasmodium falciparum. BMC Biol 11, (2013).

39. Evers, F. et al. Composition and stage dynamics of mitochondrial complexes in Plasmodium falciparum. Nat Commun 12, 3820 (2021).

40. Zhang, M. et al. Uncovering the essential genes of the human malaria parasite Plasmodium falciparum by saturation mutagenesis. Science 360, eaap7847 (2018).

41. Toenhake, C. G. et al. Chromatin Accessibility-Based Characterization of the Gene Regulatory Network Underlying Plasmodium falciparum Blood-Stage Development. Cell Host Microbe 23, 557-569.e9 (2018).

42. Chisholm, S. A. et al. The spatial proteome of the Plasmodium falciparum schizont illuminates the composition and evolutionary trajectories of its organelles. bioRxiv 2025.11.27.690946 (2025) doi:10.1101/2025.11.27.690946.

43. Tanveer, A. et al. An FtsH protease is recruited to the mitochondrion of Plasmodium falciparum. PLoS One 8, e74408 (2013).

44. Merkwirth, C. & Langer, T. Prohibitin function within mitochondria: essential roles for cell proliferation and cristae morphogenesis. Biochim Biophys Acta 1793, 27–32 (2009).

45. Shi, H., Rampello, A. J. & Glynn, S. E. Engineered AAA+ proteases reveal principles of proteolysis at the mitochondrial inner membrane. Nat Commun 7, 1–12 (2016).

46. Serricchio, M. & Bütikofer, P. A Conserved Mitochondrial Chaperone-Protease Complex Involved in Protein Homeostasis. Front Mol Biosci 8, 767088 (2021).

47. Jumper, J. et al. Highly accurate protein structure prediction with AlphaFold. Nature 596, 583–589 (2021).

48. Hallgren, J. et al. DeepTMHMM predicts alpha and beta transmembrane proteins using deep neural networks. bioRxiv 2022.04.08.487609 (2022) doi:10.1101/2022.04.08.487609.

49. Evans, R. et al. Protein complex prediction with AlphaFold-Multimer. BioRxiv https://doi.org/10.1101/2021.10.04.463034v1 (2022) xdoi:10.1101/2021.10.04.463034v1.

50. Sparkes, P. C. et al. Mitochondrial ATP synthesis is essential for efficient gametogenesis in Plasmodium falciparum. bioRxiv (2024).

51. Boucher, M. J. et al. Integrative proteomics and bioinformatic prediction enable a high-confidence apicoplast proteome in malaria parasites. PLoS Biol 16, 1–29 (2018).

52. Leonhard, K. et al. Membrane protein degradation by AAA proteases in mitochondria: Extraction of substrates from either membrane surface. Mol Cell 5, 629–638 (2000).

53. Gerdes, F., Tatsuta, T. & Langer, T. Mitochondrial AAA proteases - Towards a molecular understanding of membrane-bound proteolytic machines. Biochim Biophys Acta Mol Cell Res 1823, 49–55 (2012).

54. Amberg-Johnson, K. et al. Small molecule inhibition of apicomplexan FtsH1 disrupts plastid biogenesis in human pathogens. Elife 6, (2017).

55. Karnataki, A., Derocher, A. E., Coppens, I., Feagin, J. E. & Parsons, M. A membrane protease is targeted to the relict plastid of toxoplasma via an internal signal sequence. Traffic 8, 1543–1553 (2007).

56. Song, Z., Chen, H., Fiket, M., Alexander, C. & Chan, D. C. OPA1 processing controls mitochondrial fusion and is regulated by mRNA splicing, membrane potential, and Yme1L. Journal of Cell Biology 178, 749–755 (2007).

57. Guillery, O. et al. MetalloproteaseLmediated OPA1 processing is modulated by the mitochondrial membrane potential. Biol Cell 100, 315–325 (2008).

58. Marques, I., Dencher, N. A., Videira, A. & Krause, F. Supramolecular organization of the respiratory chain in Neurospora crassa mitochondria. Eukaryot Cell 6, 2391–2405 (2007).

59. Deitsch, K. W., Driskill, C. L. & Wellems, T. E. Transformation of malaria parasites by the spontaneous uptake and expression of DNA from human erythrocytes. Nucleic Acids Res 29, 850–853 (2001).

60. Ponnudurai, T., Lensen, A. H. W., Meis, J.F.G.M. & Meuwissen, J. H. E. Synchronization of Plasmodium falciparum gametocytes using an automated suspension culture system. Parasitology 93, 263–274 (1986).

61. Kooij, T. W. A., Rauch, M. M. & Matuschewski, K. Expansion of experimental genetics approaches for Plasmodium berghei with versatile transfection vectors. Mol Biochem Parasitol 185, 19–26 (2012).

62. Lee, M. C. S. & Fidock, D. A. CRISPR-mediated genome editing of Plasmodium falciparum malaria parasites. Genome Medicine 2014 6:8 6, 63-(2014).

63. Pasternak, M. et al. RhopH2 and RhopH3 export enables assembly of the RhopH complex on P. falciparum-infected erythrocyte membranes. Commun Biol 5, 1–12 (2022).

64. Schindelin, J. et al. Fiji - an Open Source platform for biological image analysis. Nat Methods 9, 10.1038/nmeth.2019 (2012).

65. Rappsilber, J., Ishihama, Y. & Mann, M. Stop And Go Extraction tips for matrix-assisted laser desorption/ionization, nanoelectrospray, and LC/MS sample pretreatment in proteomics. Anal Chem 75, 663–670 (2003).

66. Cox, J. & Mann, M. MaxQuant enables high peptide identification rates, individualized p.p.b.-range mass accuracies and proteome-wide protein quantification. Nat Biotechnol 26, 1367–1372 (2008).

67. PlasmoDB: An integrative database of the Plasmodium falciparum genome. Tools for accessing and analyzing finished and unfinished sequence data. The Plasmodium Genome Database Collaborative. Nucleic Acids Res 29, 66–69 (2001).

68. Tyanova, S. et al. The Perseus computational platform for comprehensive analysis of (prote)omics data. Nat Methods 13, 731–740 (2016).

69. Perez-Riverol, Y. et al. The PRIDE database resources in 2022: a hub for mass spectrometry-based proteomics evidences. Nucleic Acids Res 50, D543–D552 (2022).

70. Schalkwijk, J. et al. Antimalarial pantothenamide metabolites target acetyl-coenzyme A biosynthesis in Plasmodium falciparum. Sci Transl Med 11, (2019).

