## Supplementary material for "*Plasmodium falciparum* stomatin-like protein forms a putative complex with a metalloprotease in distinct mitochondrial loci": Response to reviewers

**Reviewer #1 (Evidence, reproducibility and clarity (Required)):**

Review of "The role of stomatin-like protein (STOML) in Plasmodium falciparum" by Verhoef et al for Review Commons.

Summary

In this study functional analysis of the stomatin-like protein (STOML) which resides in the mitochondria is performed in Plasmodium falciparum. Deletion of the stoml gene causes significant growth delay is the asexual blood stage but not in the sexual stage. The Δstoml parasites were treated to respiratory chain inhibitors to determine if they had become sensitised but there was no change suggesting STOML has another important mitochondrial function. STOML was tagged and some very high-quality imaging indicated it mainly resided at the tips of the mitochondrion. STOML main interacting partner was the protease FtsH like other organisms where modelling indicated the two proteins appear to form a vault-like macromolecular complex.

Major Comments.

While the quality of the experimental data especially the microscopy is excellent, the impact of the work is more modest considering this protein has been studied before in other organisms. The growth reduction in the Δstoml parasites was interesting but there was no robust attempt to figure out what the function of the protein was apart from one set of drug inhibition experiments. If the function of STOML could be found, then this would make the paper much more interesting. Suggested experiments are metabolomic analysis to determine if there is a block in pyrimidine synthesis which requires a functioning electron transport chain. Are there effects on mitochondrial DNA levels or organelle partitioning during cytokinesis?

We would like to thank the reviewer for the complements about the quality of the data. We agree with the reviewer that the impact of the paper could be improved if the exact molecular function of STOML at this specific mitochondrial localization can be determined. We thank the reviewer for the suggested experiments and agree that these could be valuable in further exploration of the metabolic function of STOML. We want to point out that our IFAs (Figure S3) did not show any effect of STOML KO on mitochondrial dynamics – fully matured schizonts contain normal looking, divided mitochondria – answering the question of STOML KO might affect organelle partitioning. As the PhD candidate that was driving this work has finished and moved to a different lab, we are unable to perform any large-scale experiments, such as metabolomics, proteomics, or transcriptomics. That being said, we think that the work presented in this manuscript already provides strong indications for a role of STOML in the regulation of the accessibility and therefore activity of the FtsH metalloprotease, given the strong evidence for their interaction, and the supporting structural predictions that mimic a folding as seen in a bacterial orthologue with that function. We are convinced that, within the limitations of our data which we clearly declare in the manuscript, this work still makes a valuable contribution to the field of molecular parasitology and mitochondrial biology as affirmed by reviewer 1 in their comment on the significance of this work.

Minor Comments

I am a bit confused about Fig 4 C. It looks like a heat map but is supposed to be a blue native gel. I would clarify this and at least show and example of the original gels in the Supp data. I could not find anything how the native gels were done in the Methods.

We apologise for the confusion. The data described in this panel has been reproduced from a complexome profiling data set we published previously (Evers et al 2021). We have rewritten the figure legend to clarify. Additionally, we have added a legend in the figure (figure key) indicating the relative abundance and the direction of the blue native gel to figure 4A: "*Heatmaps and line graphs based on previously published complexome profiling data showing migration patterns and relative protein abundance of STOML (red), FtsH (blue), and PF3D7_1306200 (green) in asexual blood stages (ABS) and gametocytes (GTC). Co-migration on blue native gel within the same molecular weight (MW) range (x-axis) indicates complex formation. Their relative abundance was normalized with the highest iBAQ value for a given protein set to 1 (shown red in the heatmap).*"

REFEREE CROSS-COMMENTS

I agree with the other reviewers' interpretation of the study and of additional experiments proposed.

Reviewer #1 (Significance (Required)):

As I said in Major Comments the paper is generally good and the data support the conclusions but being able to discover the function of STOML would increase the significance of the paper. The paper fills a knowledge gap particularly as to the localisation of STOML and functional importance. This paper is probably of specialised interest to basic science researchers mainly parasitologists and mitochondria specialists.

**Reviewer #2 (Evidence, reproducibility and clarity (Required)):**

This work sheds new light on mitochondrial biology of the human infecting malaria parasite, Plasmodium falciparum. The study investigates the essentiality and function of a stomatin-like protein (STOML) in asexual blood stages. The P. falciparum STOML protein is a member of the highly conserved SPFH group of four proteins in P. falciparum that localize to the mitochondria. Previous studies in the murine malaria parasites, P. berghei, have shown SPFH proteins are critical for Oocyst development. However the importance of STOML in asexual blood stages remains unclear as clonal knockout lines could not be obtained, suggesting a critical role for this protein in parasite development. In this study, the authors successfully generated a P. falciparum knock-out line by replacing the STOML coding sequence with GFP. An elegant competition growth assay was then used to show that the asexual blood stages of the knockout line exhibit a modest growth defect compared to a control population (expressing mScarlet) in culture. This growth phenotype was not associated with obvious changes in mitochondrial morphology or a defect in respiratory chain activity as shown by ultrastructural changes or increased sensitivity to respiratory chain inhibitors. Moreover, the STOML knockout was able to differentiate to gametocytes which have larger mitochondrion and more active TCA cycle, suggesting that STOML is not critical for these changes. Analysis of STOML localization throughout asexual blood stage development using live microscopy indicated that the STOML-mNEON protein was localized to punctate structures that appear to coincide with mitochondrial branching points; suggesting a possible role in mitochondrial segregation. The authors also show that STOML migrates at a large 2MDa complex via BN-PAGE and interacts with FtsH and a novel protein of unknown function (which intriguingly was predicted to localize to the apicoplast). Based on these different lines of evidence the authors propose a model in which STOML forms a 22-multimer barrel similar to bacterial family members that complex with FtsH to possibly regulate the accessibility and proteolytic function of FtsH. Overall, the data reported in this study are clear and support the conclusions, and add to the growing body of work on the role of STOML proteins in different protists.

We would like to thank the reviewer for their kind words on our work.

Major points :

It is possible that tagging of STOML with mNEON green may affect both the localization and oligomerization of this protein in the inner mitochondrial membrane. It would be useful to additionally validate the localisation of HA only tagged forms to confirm the punctate pattern detected.

We agree with the reviewer that the large fluorescent mNeonGreen tag might affect localization or oligomerization of STOML and that it would therefore be useful to confirm STOML localization in the *stoml-HA* parasite line. We have therefore performed an immunofluorescence assay (IFA), which confirmed localization of *Pf*STOML-3HA to mitochondrial branching points and branch endings, consistent with the localization observed with live imaging of *Pf*STOML-3HA-NG (Figure S1D). We have added the following text to the result section of the manuscript:

Line 245: “An immunofluorescence assay confirmed that *Pf*STOML-3HA localizes to mitochondrial branching points and branch endings, consistent with the pattern observed in live imaging of *Pf*STOML-3HA–mNG (Figure S1D, 2). This demonstrates that the addition of the large mNeonGreen tag does not affect the native mitochondrial localization of *Pf*STOML.”

Reviewer #2 (Significance (Required)):
Molecular processes involved in the biogenesis and functioning of the mitochondria in Plasmodium parasites are of considerable interest as potential drug targets. Previous work investigating the role of the SPFH proteins in Plasmodium species have given incomplete or conflicting results. This comprehensive study investigates the role of the STOML protein, a member of the SPFH family in human infective P. falciparum asexual blood stages and gametocytes. Using a range of genetic and molecular techniques this study shows that PfSTOML integrats into large protein complex in the mitochondria and has a potential role in regulating mitochondrial dynamics, although loss of this protein only has a modest affect on parasite growth in RBC in vitro. The findings will be of broad interest to researchers in the parasitology and mitochondrial fields.

Reviewer #3 (Evidence, reproducibility and clarity (Required)):

Summary:
The authors study an understudied mitochondrial SPFH protein, called stomatin-like protein (STOML), in the important human parasite Plasmodium falciparum. Many recent studies, including by the authors, have shown Plasmodium mitochondria to have many unique features and biology compared to that of the host. However, their mitochondria remain relatively understudied and has great potential for therapeutic intervention, making studying their biology both timely and important. Using a variety of molecular parasitology approaches, via genetic manipulation of parasites, microscopy, proteomics and structural modelling the authors to probe the role and mark unique features of this protein in Plasmodium falciparum.

We thank the reviewer for taking the time and effort to look at the manuscript.

Major comments:

1) Involvement with respiratory chain:

Lines 174-179. Fig S4: This analysis would be strengthened by including a positive control. I appreciate that sensitivity to respiratory chain inhibitors has previously been demonstrated (on a separate note, a reference to one of these previous studies should be included and the rationale for the experiment spelt out for readers less familiar with the field), but without inclusion of a positive control (e.g. a parasite line known to be defective in the respiratory chain) the data presented here are negative data without a positive control from the same experiment. This analysis would also be strengthened by inclusion of another replicate and noting the IC50s of these drugs and if these are in agreement with the published literature.

We apologise for omitting to include a reference to previous studies and have now included three: Sheokand et al (2024), Pietsch et al (2023), and Narwal et al (2023). These studies show that deletion of *Pf*Rieske (subunit of complex III of the respiratory chain), *Pf*PNPLA2 (patatin-like phospholipase), and *Pf*OPA3 (OPA3- like domain-containing protein), respectively, affect the sensitivity to drugs targeting the respiratory chain. These three studies have robustly demonstrated that this assay can give an indication of the integrity of the respiratory chain. However, we respectfully disagree with a need for a "positive" control. The effectivity of these drugs has been very robustly demonstrated in many papers and by testing the drugs on WT parasites we have included a positive control for drug activity. Of note, all the $IC_{50}$s observed in our study correspond to previously determined values. If the STOML KO line would have been more (or even less) sensitive than WT, this would most certainly have been obvious from the data. We do agree that a lack of shift in sensitivity to these drugs does not necessarily rule out a function of STOML in the respiratory chain assembly. However, it is a clear indication that it is likely not (supported by the fact that they can make healthy gametocytes and gametes). We chose our wording carefully to reflect this. To further strengthen the data, we have now combined all drug assays that we performed instead of showing one representative experiment. We have included the IC50 values with their 95% confidence interval below each graph, which are similar to previously reported IC50 values. For most drugs, we have four independent biological replicates, except for chloroquine, proguanil and MMV183, for which we have two independent replicates, which has been clearly indicated in the figure legend.

The conclusion of no defect in the respiratory chain would also be strengthened by additional experiments directly measuring respiratory activity pointing to the same conclusion (mitochondrial morphology displayed in S3 and S5 are fairly indirect and can be affected by numerous other processes). Examples include extracellular flux analysis (PMID:37969754), CIII assay (PMID: 29775797), Mitotracker (combined with microscopy analysis) assessment of mitochondrial membrane potential (PMID: 34612205; 37126705). It may be that these assays are more sensitive than the change in susceptibility to inhibitors, and that by only assaying respiratory chain function through a single experiment a mutant phenotype might be missed. I mark these additional experiments as OPTIONAL as I appreciate it represents a significant amount of work, but it would add significantly to the conclusion of an unaffected respiratory chain.
We agree with the reviewer that additional experiments as suggested above would strengthen our conclusion of *Pf*STOML not being involved in respiratory chain assembly. However, as indicated above the PhD candidate leading this work has left the lab and we are unfortunately not in the position to perform additional experiments that cost significant amounts of time and/or resources. As written in a reaction to the previous comment, several papers have shown involvement of proteins in the function of the respiratory chain can be shown using this drug assay. This, together with the fact that the *stoml*$^{-}$ line can make healthy gametocytes with normal-looking mitochondria, argues against *Pf*STOML involvement in the respiratory chain.

Overall, given the human STOML homolog appears to play a role in the assembly of the respiratory chain, the fact the Plasmodium protein does not seems one of the major findings of the paper, and so additional evidence (along with extra repeats and references to previous literature in the field) would significantly strengthen this section.

2) A major finding of this paper is the interaction of STOML with the zinc-dependent metalloprotease FtsH. However, the presentation of the proteomic evidence for this lacks some details.

-The authors say "we performed two independent pulldown experiments": details are lacking, both in text and materials and methods, about whether biological replicates were performed within each experiment or are single replicate experiments. The findings of interaction would of course be much stronger if multiple replicates have been performed.
We would like to further clarify the setup of the pulldown experiments. As stated in the text line 235-243, two completely independent pulldown experiments were performed. In the first experiment, the *stoml-NG* parasite line was used and the pulldowns were performed with anti-mNG coated beads and empty binding control beads. In the second experiment, *stoml-HA* parasite line was used and the pulldowns were performed with anti-HA coated beads and empty binding control beads. In the materials and methods (line 541 onwards) we describe the setup of both pulldown experiments. For each

experiment, an input of 18x 30-ml culture synchronized parasite pellets were used, which we harvested from 18 different parasite cultures over a period of several months. After obtaining the organellar fraction through nitrogen cavitation, pulldowns with magnetic beads were performed in three technical replicates. To clarify this, we added the following sentences to the materials and methods (line 548): "*Both pulldowns were carried out with three technical replicates*". Even though the two independent experiments were done with different protocols on two different genetic backgrounds, we identified STOML and FtsH in both – providing compelling evidence that these are true interactors. This conclusion is corroborated by published work, including our own complexome profiling data and existing literature about STOML-FtsH interactions in other species. Consequently, we think it is wasteful to perform a third highly time intensive and costly experiment.

- Is there a data availability statement: the raw data would ideally be deposited in an appropriate repository (e.g. PRIDE).

We thank the reviewer this comment. The mass spectrometry data has been uploaded the PRIDE database and will be made available upon publication. A password to enable editors and reviewers to access the data will be provided directly to the editor. To clarify we have added the following sentence to the materials and methods (line 576): "*The mass spectrometry proteomics data have been deposited to the ProteomeXchange Consortium via the PRIDE partner repository with the dataset identifier PXD039772.*"

-The supplemental data should include data for all detected proteins rather than just those found enriched (additionally column headings should contain more details: what does "mito score" refer to?; why have t-test value but not fold change information been included?). This would allow readers to tell, for example, if the 119 proteins detected as enriched in experiment 2 (but not 1) were a) detected, but not enriched, in the first experiment, or b) not detected at all.

We thank the reviewer for this comment and agree that including all detected proteins in the supplemental data, and not just the significant hits could be useful for the reader. Therefore, we have now updated the supplementary table S2 and S3 accordingly. We have added the following clarification to the legends of these tables: "*This table includes all proteins detected in the STOML-3HA-NG pulldown, indicated by their gene ID in the first column and the annotation in the second column. T-test significance, unique peptides, log p-value and t-test difference are indicated in the following columns. Mitochondrial prediction score is indicated in the last column as "Mito score", which refers to the ranking of the protein in the mitochondrial proteome as shown by Esveld* et *al.*"

-From previous complexome analysis the authors show that STOML and FtsH partially co-migrate in gametocytes but not in ABS. Can the author's comment which stage the IP most corresponds to: if it is ABS can the authors comment about how the proteins apparently interact (from IP evidence) but do not co-migrate (complexome evidence).

The life-cycle stages analyzed in the CoIPs were late-stage ABS parasites (stated in materials and methods line 534) and consequently most closely resemble the mixed ABS analyzed in the complexome profiling.

Despite our attempts to keep detergent, enrichment and cell lysis conditions similar for both approaches, there are some inherent differences between methods that could underlie differing outcomes.
First of all, the Coomassie brilliant blue dye used in complexome profiling to give charge for more effective electrophoresis, has been shown to negatively affect some protein interactions and long time under an electric field with different buffer from CoIP can lead to further dissociation (sources for CBB destabilizing effect PMIDS: 16287510, 16220535, 23744064).
Second, the concentration and immobilization of STOML on beads during CoIP effectively increases local concentration of the bait protein which increases binding and rebinding of low affinity interactors. Complexome profiling has no such built-in concentration mechanism and disassociations that occur during electrophoresis or initial solubilization are more likely to remain disassociated. The same holds true for potential bridging factors such as lipids, proteins or nucleic acids that could be required for the interaction between STOML and FtsH.
Finally, the concentration and isolation that occurs during CoIP also facilitates easier identification through mass spectrometry, which is important for low abundance targets such as FtsH where this lowered sensitivity can lead to artificially spiky and noisy detection, that can further differ between samples even if abundances are similar. If for example in a given fraction of the gel there is the same amount FtsH in both ABS and gametocytes but the ABS sample has more competing proteins with peptides that have a similar mass over charge ratio compared to those generated by FtsH, it will be less likely to be detected in the ABS sample, despite same abundances.
A combination of these factors could explain both the less compelling evidence of complexome profiling and the differences between ABS and gametocytes data.

Another factor that sets the two methods apart, is the type of data that they produce.
The migration profiles resulting from complexome profiling represent all (sub)assemblies of a given protein. The subset of STOML that binds FtsH could be a minority peak and consequently looking at the relative migration pattern, might not show strong comigration despite a putative real interaction between STOML and FtsH. The broad migration pattern of STOML is indicative of either a promiscous protein with many (self) interaction states or alternatively a disintegrating large protein complex with a smear of breakdown products in line with the earlier described disassociation risk in complexome profiling.

Taking this factor into account, we can look at intensity based absolute quantification (iBAQ) values rather than only relative migration patterns.
In line with only a minority fraction of STOML associating with FtsH, we see that STOML is much more prevalent than FtsH by a factor of 173x and 130x in gametocytes and ABS respectively. Plotting these values we can see that across both gametocytes and ABS, STOML and FtsH cooccur in the same fractions but that the broad migration pattern and majority of STOML protein that does not appear to associate with FtsH skews the picture when plotting relative migration pattern.

To clarify this, we have added Figure S6 to the manuscript, which contains of two graphs showing the absolute protein values (iBAQ) of STOML, FtsH and PF3D7_1306200 in ABS

and gametocytes. We have also added the following text to the corresponding result paragraph (line 260):
*"Both in ABS and gametocytes, FtsH runs at ~2.5 MDa, which in gametocytes corresponds with the most abundant STOML peak (Figure 3A, S6)*[38]*. In ABS, FtsH appears to not perfectly comigrate with STOML when looking at relative migration patterns. However, taking absolute protein quantity (iBAQ value) into account, the FtsH migration peak overlaps with the broad STOML migration peak (Figure S6). STOML is much more prevalent than FtsH by a factor of 173x and 130x in gametocytes and ABS respectively, which could explain why they do not show strong comigration in the normalized graph, but do have overlapping migration pattern when looking at absolute values. The broad migration pattern of STOML may represent different (sub)assemblies of the protein complex. Perhaps, only a subset of STOML forms a complex with FtsH."*

So taken together the complexomics data across both gametocyes and ABS, while not providing clarity on interaction on their own, are fully compatible with true and specific interactions captured by CoIP.

-OPTIONAL: validating this interaction through further experiments would strengthen this conclusion, for example introducing a second epitope tag to FtsH and performing co-ip and immunoblot experiments, or tagging FtsH and performing an IP and, through a similar proteomic experiment, seeing if STOML is detected.
Although we agree with the reviewer that STOML-FtsH interaction would be further confirmed with a reciprocal IP, we do not think these experiments will be possible. Other groups have been unsuccessful in tagging of FtsH (Amberg-Johnson 2017). Furthermore, we think the consistent pulldown of FtsH with two independently tagged STOML lines is strong evidence that FtsH is indeed an interaction partner of STOML. Additionally, the interaction of SPFH proteins with metalloproteases is highly conserved among prokaryotes and eukaryotes. The human, *Trypanosoma brucei,* and *Neurospora crassa* homologs of *Pf*STOML form large complexes with YME1L/FtsH, which are FtsH homologs. This indicates that FtsH is a very likely interaction partner of STOML, since the STOML-FtsH complex is highly conserved across different eukaryotes (explained in line 410 - 420).

Overall, despite being a major finding of the paper the description of the experiments and data availability are not quite sufficient to assess the conclusions. This can be easily remedied through text changes and addition of more extensive supplemental data tables.

Minor comments:

1) Title: the title talks about the "role" of STOML in Plasmodium falciparum but I feel at the end of this study the role is still unknown (which the authors acknowledge Line 427-428). The authors convincingly show it is important and that it has some interesting features (e.g. punctate localization to branching points of mitochondrial network) but the exact role it plays will require further study. The authors should consider changing the title to reflect this.

We have changed the title to: *Plasmodium falciparum* stomatin-like protein forms a putative complex with a metalloprotease in distinct mitochondrial loci

2) Line 71- 2: "as well as an unusual myzozoan-specific prohibitin-like protein (PHBL)"- it may make it easier for a general reader to introduce/ define the term myzozoan.
We have changed to:
"PHBL is specific to the unicellular Myzozoa, a clade that includes apicomplexan parasites and free-living dinoflagellates."

3) Figure S1A contains a schematic for tagging the gene on the left, and then knocking the gene out on the right. Given that the KO is described first in the first results section (starting line 109), and the tagging described after that in the third results section (line 194 onwards) it would aide readability to have the diagrams in the same order as the text, and perhaps separate them into separate panels. In fact, epitope tagging may be better with its own supplemental figure later in the order.
We thank the reviewer for their comment, and we have changed this figure accordingly.

Also, for figure S1A- schematic should label "3-HA" instead of "HA" to match text description. S1B,C- label on figure "stoml-NG" should say "stoml-NG-3HA" to match text descriptions. Inconsistency in how this line is referred to.
We have changed the label in Figure S1A to 3HA in the figure to match text description.
We are consistently describing the parasite line which contains STOML tagged with 3HA-NG as *stoml-NG*, while *Pf*STOML-3HA-NG is used to describe the expressed STOML protein which is tagged with 3HA-NG. Therefore, we will not change this in the label of the figure to avoid inconsistency.

4) Figure S2 has a schematic in a showing the integration of an mScarlet repair plasmid. Then in B there is the PCR integration for "cyto-mScarlet" but also "cyto-mNG"- is there a missing schematic for creating the "cyto-mNG" line?
For generation of the cyto-mScarlet and cyto-mNG lines, the same strategy is used (except one has an mScarlet and the other an mNG protein). To clarify this, we have added made the gene encoding for the fluorescent protein (mScarlet or mNG) yellow in figure S2 and changed the text in the legend: "*Schematic overview of transfection strategy <u>to generate</u> <u>cyto-mScarlet</u> <u>and</u> <u>cyto-mNG</u>*." And "*DNA breaks are repaired by double homologous recombination with a repair plasmid containing 5' and 3' homology regions (HRs) and a <u>fluorescent protein (FP, mScarlet or mNeonGreen</u>) under the control of the H2B promotor and PBANKA_142660 bidirectional 3'UTR*.

5) When describing the mitochondrial morphology experiments (line 167-173) it would be good to reference previous work that shows that defects in mitochondrial function can lead to defects in mitochondrial morphology. This would provide a much clearer rationale for why these experiments were performed.
The rationale to perform this experiment is to study if knockout of STOML alters mitochondrial dynamics, and not necessarily to study the effect on mitochondrial function. The localization of STOML to specific foci on the endings and branching points of the mitochondrial network suggest a possible function of STOML in mitochondrial

division or segregation. E.g. *Pf*DYN2 (dynamin 2) has been shown to be involved in mitochondrial and apicoplast division, localizing to mitochondrial branching points. Disruption of PfDYN2 significantly altered mitochondrial morphology in schizonts (Morano 2025). STOML KO did not cause any obvious differences in mitochondrial morphology and there were no clear mitochondrial division or segregation defects observed in these parasites, suggesting that STOML does not play an essential role in these processes. This is explained in the discussion line 351 – 354.

6) lines 194-217: Could the authors clarify if they mean the protein is in the mitochondria (does the signal overlap) or that they see the signal beside the mitochondria (i.e. in close proximity but not actually in the mitochondria).

The STOML-3HA-NG signal is largely (but not completely) overlapping with the mito-mScarlet signal, as can be observed from the 3D visualizations in Figure 2B. This suggest that STOML resides at the mitochondrial membrane, and not the mitochondrial matrix, while we lack the resolution to distinguish between the inner mitochondrial membrane or outer mitochondrial membrane. However, STOML is a highly conserved proteins, and homologs in human (Wai 2016), *Trypanosoma brucei* (Serricchio 2021) and *Neurospora crassa* (Marques 2007) have shown that STOML resides in a large protein complex at the inner mitochondrial membrane. Therefore, it is highly likely that STOML in *P. falciparum* also localizes to the inner mitochondrial membrane, especially given that it forms a complex with FtsH, an i-AAA protease which are also known localize to the IMM. To clarify, we added the following sentence to the text in line 211: *The PfSTOML-3HA-NG foci are largely, but not completely overlapping with the mito-mScarlet signal in trophozoite and schizont stages (Figure 2B), suggesting localization to the mitochondrial membrane, although we lack the resolution to distinguish between IMM and OMM.*

7) Line 236: figure 4C is referred to before figures 4A-B. It is more customary for the order in the text to reflect the panel order. Labels in 4C- "GTS" not defined.

We have corrected the panel order in Figure 4 accordingly. We have also clarified GTS in the figure legend.

8) Reference 26 appears to have a typo in it.

Corrected

9) Line 89: missing reference for PbSTOML localization.

Changed accordingly

Reviewer #3 (Significance (Required)):

This study tackles an understudied protein of Plasmodium falciparum mitochondria and therefore provides interesting new data about mitochondrial biology in this parasite. As noted above, the "role" of this protein is still unknown, but this study provides a number of interesting initial experimental observations, providing a platform for future work to elucidate its function.

A greater advance in understanding may have been obtained had the authors performed more in-depth follow-on experiments about any one of these observations. For example,

the authors observe a unique localization pattern, but do not probe this with further experimentation and explore how this may be achieved and how it contributes to function.
This study has the potential to be of interest to a wide-range of basic researchers in both the parasitology and mitochondrial biology fields.

Expertise: molecular parasitology, mitochondrial biology.
