## Supplemental information for "*Plasmodium falciparum* stomatin-like protein forms a putative complex with a metalloprotease in distinct mitochondrial loci"

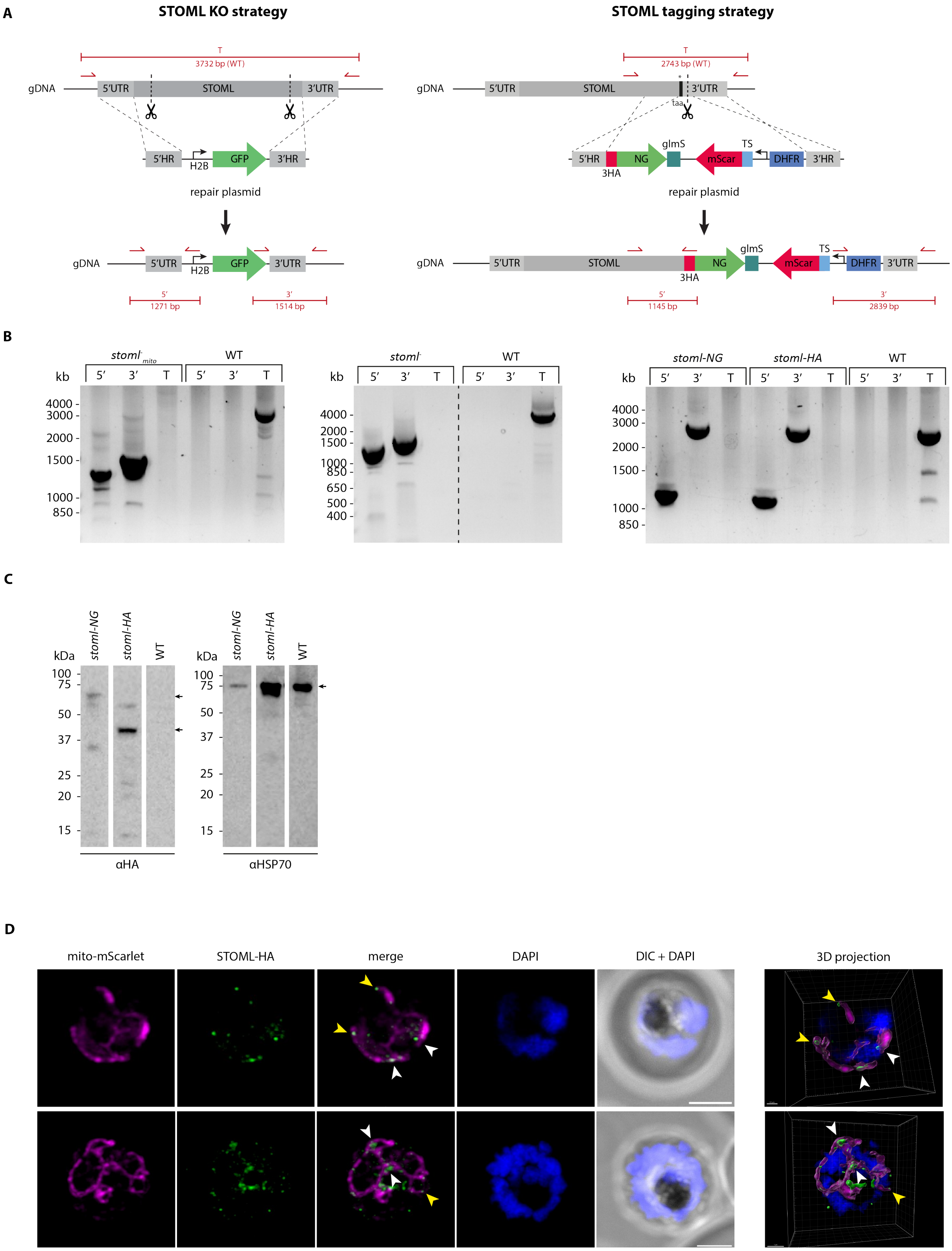


***Figure S1. Generation of* STOML *tagging and KO parasite lines.*** *A) Schematic overview of* STOML *tagging and KO strategy. For tagging of* STOML *with 3HA-NG-glmS or 3HA-glmS, CRISPR-Cas9 (indicated by scissors) is used to introduce a double-strand break to facilitate integration of the linear repair constructs 3HA(-NG)-glmS tag directly after* STOML *before the stop codon, while at the same time integrating a mito-mScarlet mitochondrial marker and a DHFR drug selection cassette. For STOML KO, two CRISPR-Cas9 introduced DNA breaks at the 5' and 3' of the gene will be repaired by the linearized HDR plasmid. After integration,* STOML *will be replaced by GFP under the control of the H2B promotor. B) Diagnostic PCR of* stoml-NG, stoml-HA *and* stoml$^{-}_{(mito)}$ *parasite lines with integration specific primer combinations (indicated in panel A), demonstrating successful 5' and 3' integration and the absence of WT parasites (T=total). C) Western blot analysis showing expression of STOML-3HA-NG (73 kDa) and STOML-3HA (47 kDa) at expected sizes using anti-HA antibody and anti-HSP70 for loading control. D) Fluorescence microscopy of* stoml-HA *with anti-HA antibody (green), mito-mScarlet mitochondrial marker (magenta), DAPI for DNA visualization (blue), and DIC. Images are maximum intensity projections of Z-stack confocal Airyscan images, except images in right panel, which are 3D visualizations generated with Imaris analysis software. Arrowheads indicate* Pf*STOML-3HA signal at mitochondrial branching points (white) or mitochondrial branch endings (yellow). Scale bars are 2 µm for maximum intensity projections and 1 µm for 3D visualizations.*

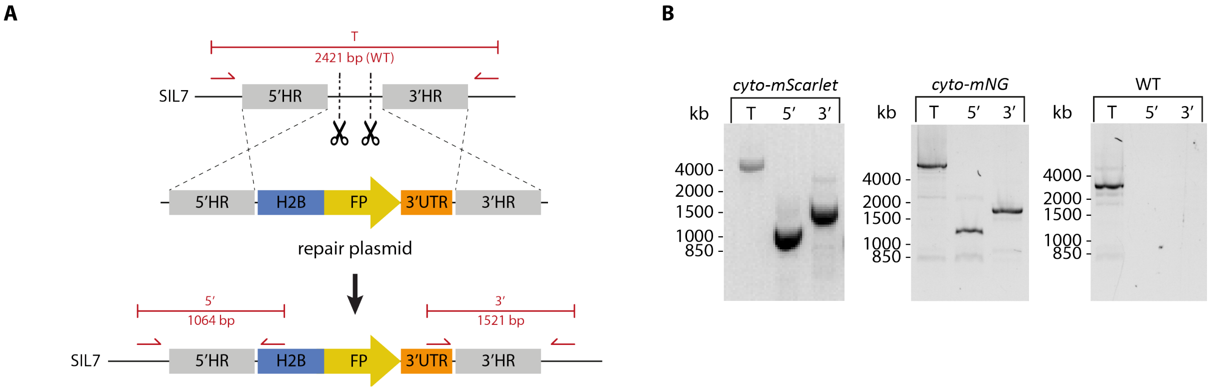


***Figure S2. Generation of*** **cyto-mScarlet** ***and*** **cyto-mNG** ***parasite lines.*** *A) Schematic overview of transfection strategy to generate cyto-mScarlet and cyto-mNG. CRISPR-Cas9 and two guides were used to generate double stranded breaks in a silent intergenic locus (SIL7), characterized in Verhoef* et al.[1] *(indicated by scissors). DNA breaks are repaired by double homologous recombination with a repair plasmid containing 5' and 3' homology regions (HRs) and a fluorescent protein (FP, mScarlet or mNeonGreen) under the control of the H2B promotor and PBANKA_142660 bidirectional 3'UTR. B) Diagnostic PCR of* cyto-mScarlet *and* cyto-mNG *parasite lines with integration-specific primer combinations (indicated in panel A), demonstrating successful 5' and 3' integration and the absence of WT parasites (T=total).*

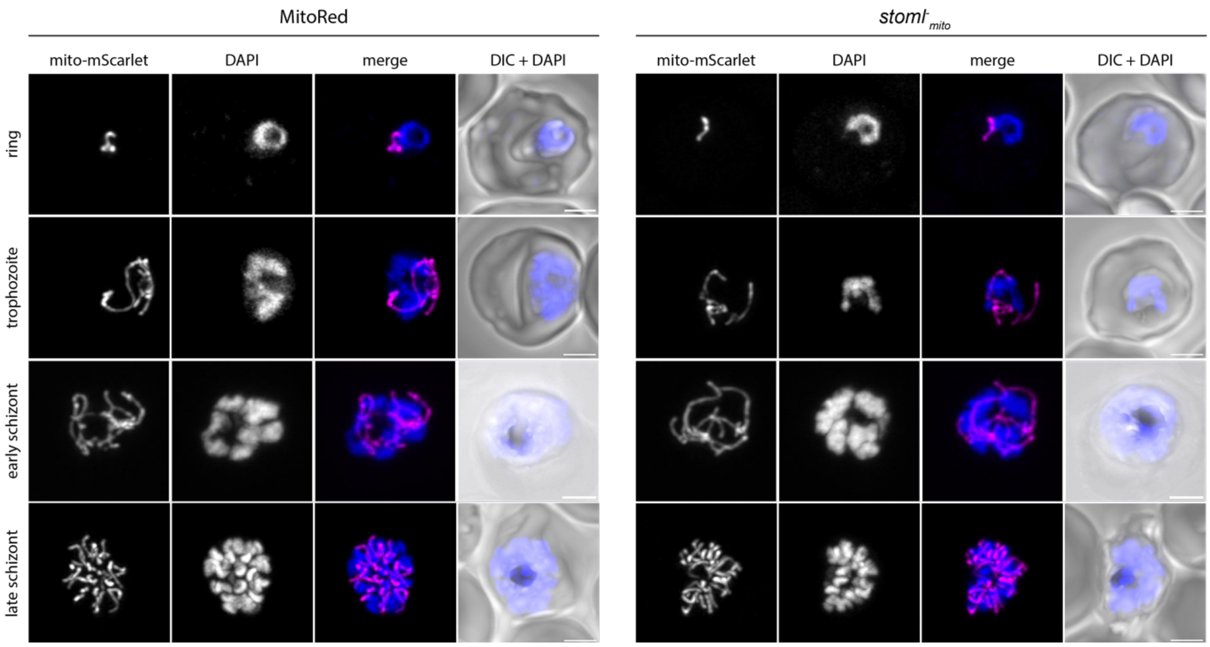


***Figure S3. Mitochondrial morphology in stoml$^{-}_{mito}$ ABS parasites.*** *Fluorescent microscopy of stoml$^{-}_{mito}$ and MitoRed (WT) parasites during ring, trophozoite, early and late schizont stages. The mito-mScarlet signal is preserved after fixation and can be observed without antibody staining. DNA was stained using DAPI. Images are maximum intensity projections of Z-stack confocal Airyscan images. Scale bars, 2 $\mu$m.*

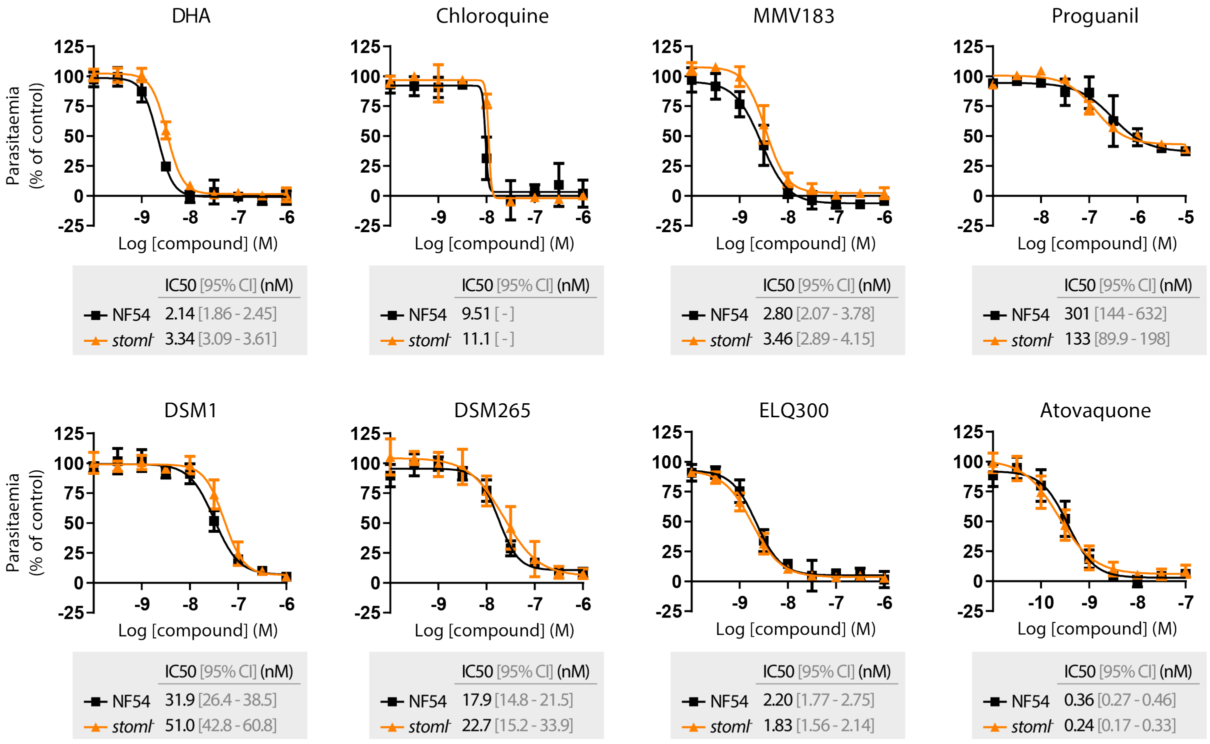


***Figure S4. Sensitivity of*** **stoml$^{-}$** ***parasites to anti-malarial compounds.*** *Drug sensitivity assay for* P. falciparum *NF54 and* stoml$^{-}$ *parasites. The graphs show average values for mean parasite density relative to controls for asexual blood-stage replication assay and represent one of the two independent replicates. Error bars indicate SEM determined from two technical replicates per experiment. The data were analyzed using nonlinear regression in GraphPad Prism. Proguanil, DSM1, DSM265, ELQ300, and Atovaquone are compounds targeting the parasite mitochondrion, while DHA, chloroquine, and MMV183 are non-mitochondrial compounds.*

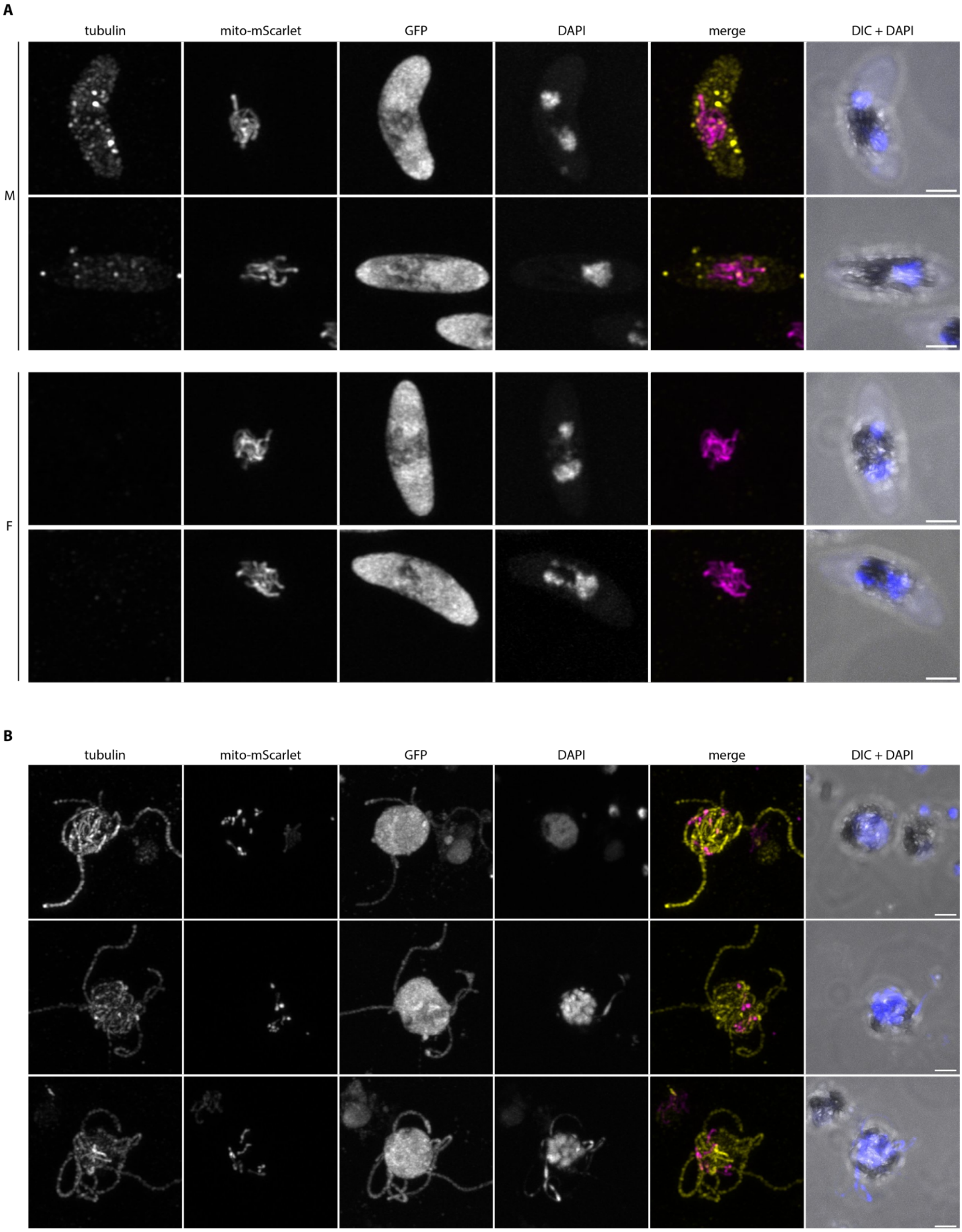


***Figure S5.*** **Stoml$^{-}_{mito}$** ***parasites develop to healthy gametocytes that exflagellate.*** *A) Fluorescent microscopy on male (M) and female (F)* stoml$^{-}_{mito}$ *stage V gametocytes. Parasites were stained for tubulin (yellow) to distinguish male (high $\alpha$-tubulin signal) from female (low $\alpha$-tubulin signal) gametocytes. B) fluorescent microscopy on exflagellating* stoml$^{-}_{mito}$ *male gametes at 20 minutes after activation. Parasites were stained with tubulin to visualize axonemes. A-B) Visualization of mito-mScarlet mitochondrial marker (magenta), cytosolic GFP, DAPI for DNA visualization (blue), and DIC. Images are maximum intensity projections of Z-stack confocal Airyscan images. Scale bars, 2 $\mu$m.*

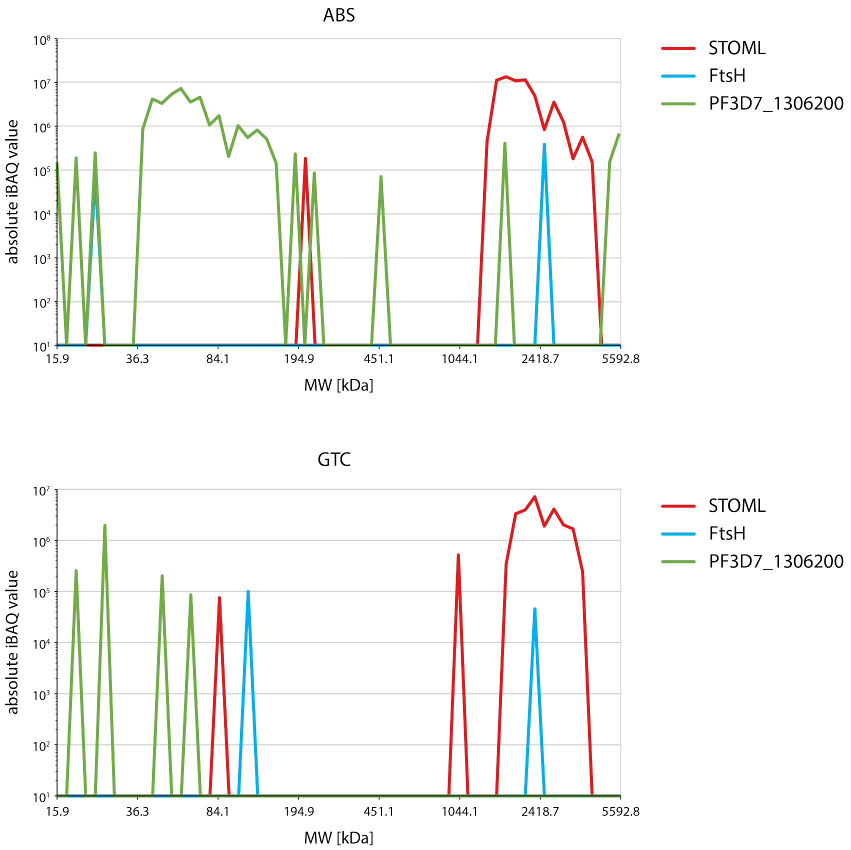


***Figure S6. Absolute protein abundance of STOML, FtsH and PF3D7_1306200 across protein complex migration on native PAGE gel.*** *Line graphs based on previously published complexome profiling data[2] showing migration patterns and absolute protein abundance (iBAQ value, logarithmic scale) of STOML (red), FtsH (blue), and PF3D7_1306200 (green) in asexual blood stages (ABS) and gametocytes (GTC). Co-migration on blue native gel within the same molecular weight (MW) range (x-axis) indicates complex formation.*

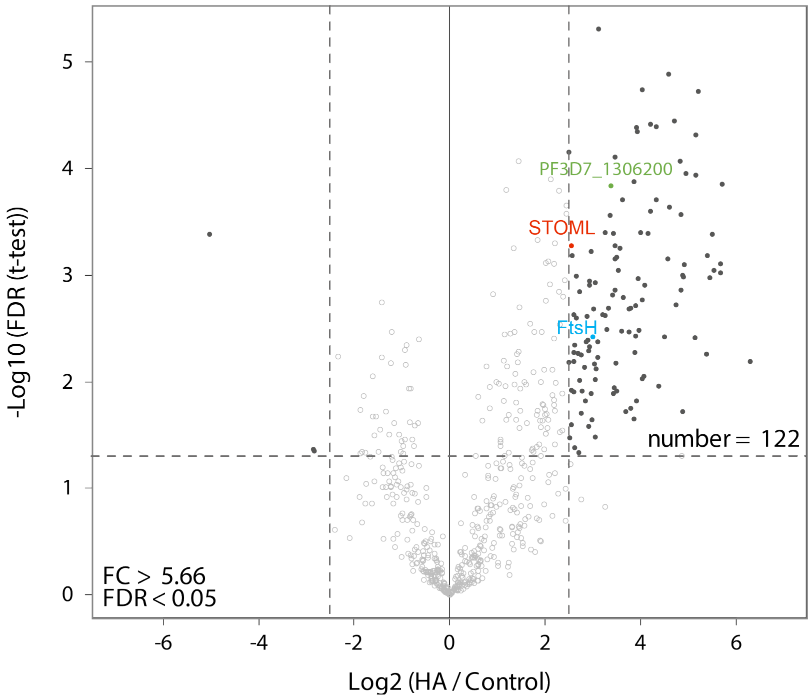


***Figure S7. Identification of STOML interacting proteins with co-immunoprecipitation.*** *Anti-HA immunoprecipitation of* Pf*STOML-HA containing complexes. The volcano plot showing mean* $log_2$ *fold changes (FC) and* $-log_{10}$ *false discovery rate (FDR) for anti-HA pulldown in comparison with control pulldown. Horizontal and vertical dotted lines indicate* $log_2$ *FC > 2.5 (FC > 5.66) and* $-log_{10}$ *FDR > 1.301 (FDR > 0.05) respectively. Dark dots represent proteins that are highly enriched or reduced in the anti-HA pulldown compared to the control pulldown.*

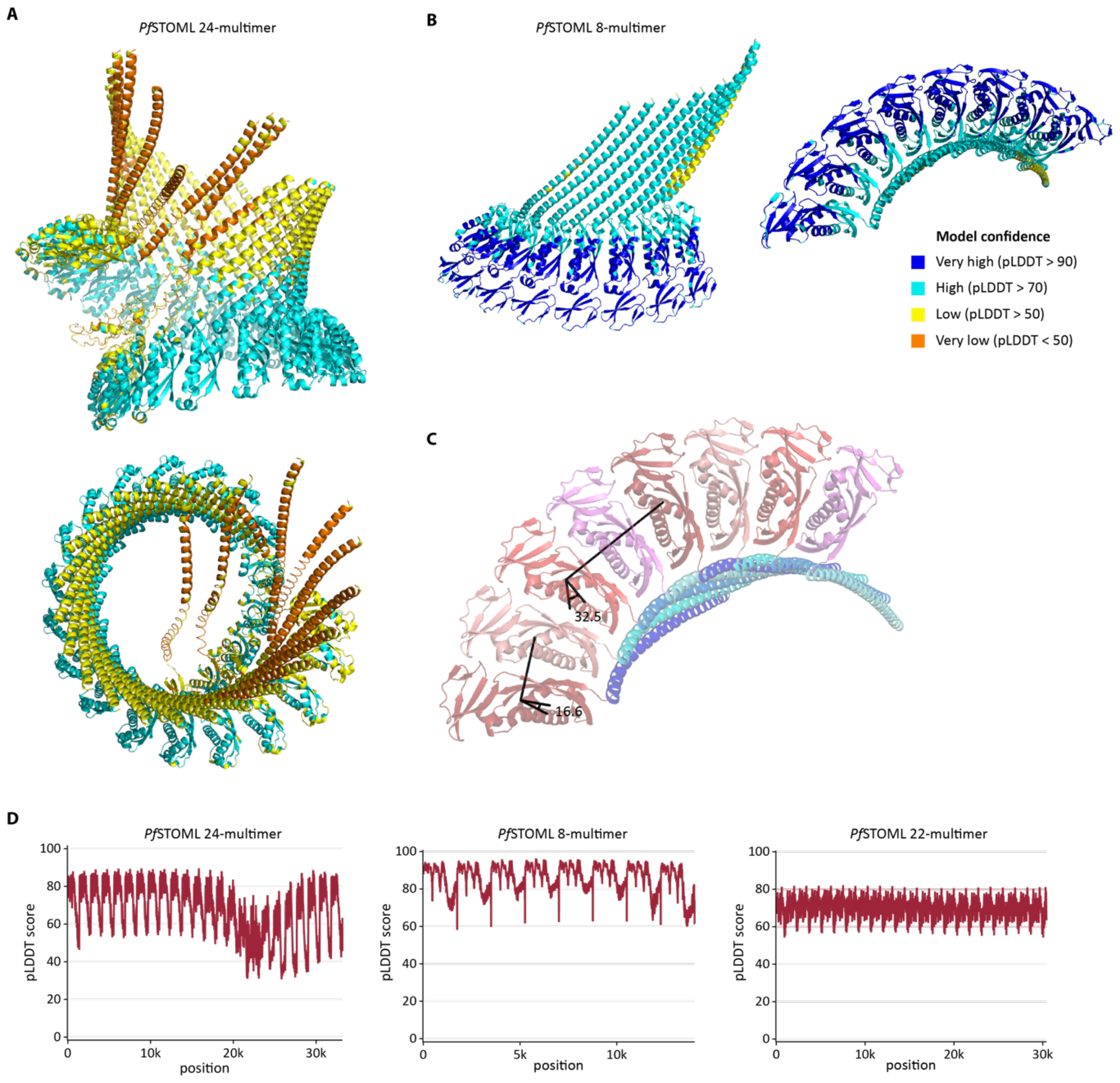


***Figure S8. AlphaFold2 structure prediction of* Pf*STOML multimers.*** *A) AlphaFold2 prediction of* Pf*STOML 24-multimer with side view (top) and top view (bottom). B) AlphaFold2 prediction of* Pf*STOML 8-multimer with side view (left) and top view (right). Coloring in A and B represent model confidence as indicated by the color legend in B. C) Top view of predicted* Pf*STOML 8-multimer structure, indicating the angles measured between SPFH domains of different STOML proteins in the complex. D) Graphs with pLDDT scores representing model confidence of predicted* PfSTOML *24, 8, and 22 multimer structures.*

### Table S1. Primer and guide RNA sequences for generation of repair and guide plasmids.

| Primer name | Primer function | Sequence | Restriction site |
|---|---|---|---|
| STOML KO repair plasmid | | | |
| JV047 | STOML KO 5'HR F | AAATATCCCGGGCCTGCTATAGATTAATTGTGTCCC | XmaI |
| JV048 | STOML KO 5'HR R | TAATATCTCGAGGTTCATCTTCTCATGAATTTTTATCAATC | XhoI |
| JV049 | STOML KO 3'HR F | AATATACCATGGGAACAACTAATTTGATTCAAGTTTTGC | NcoI |
| JV050 | STOML KO 3'HR R | ATTAATGAATTCAATATACACAACTTATTTTTTATGTTTTCCC | EcoRI |
| STOML tagging repair plasmids | | | |
| JV084 | STOML tag 5'HR F | AATTAAGGTACCGGAACCATTTAGGTTTTGTGATTATACC | KpnI |
| JV113 | STOML tag shield R | CTCTTTCTCCTTCGCTCTGTAAAATTTCAGC | |
| JV114 | STOML tag shield F | GCTGAAATTTTACAGAGCGAAGGAGAAAGAG | |
| JV085 | STOML tag 5'HR R | AATAATGGATCCGTTGTTCATATCAGAATGAATTTGTTTTTG | BamHI |
| JV027 | STOML tag 3'HR F | ATTATTGAATTCATATGAAAACAACTATATTTTTATGAAAGC | EcoRI |
| JV086 | STOML tag 3'HR R | TTAAAAGCCGGCTATAAATTGTACGTGTATTATTCATTACTC | NgoMIV |
| Cyto-mScarlet and cyto-mNG repair plasmids | | | |
| JV064 | mScarlet F | AATAAAGCTAGCATGGTGAGCAAGGGCGAGG | NheI |
| JV072 | mScarlet R | AATAATCTTAAGTTACTTGTACAGCTCGTCCATGC | AflII |
| JV062 | mNG F | AATAAAGCTAGCATGGTGAGCAAGGGCGAGGAG | NheI |
| JV072 | mNG R | AATAATCTTAAGTTACTTGTACAGCTCGTCCATGC | AflII |
| Guide RNA sequences | | | |
| JV051 | STOML KO G1 F | TATTGATGTATGATACCCTTTAGCT | |
| JV052 | STOML KO G1 R | AAACAGCTAAAGGGTATCATACATC | |
| JV053 | STOML KO/Tag G2 F | TATTGGCTGAAATTTTACAAAGTGA | |
| JV054 | STOML KO/Tag G2 R | AAACTCACTTTGTAAAATTTCAGCC | |
| Pf0084 | SIL7 guide 1 sense | TATTGTATATGTGGTAATAAATAAA | |
| Pf0085 | SIL7 guide 1 antisense | AAACTTTATTTATTACCACATATAC | |
| Pf0086 | SIL7 guide 2 sense | TATTGATTCAATATAATAAGGTCAA | |
| Pf0087 | SIL7 guide 2 antisense | AAACTTGACCTTATTATATTGAATC | |
| Integration PCRs | | | |
| JV108 | Int PCR STOML tag 5' F | AATATGGAACCGAGCTAAAGGG | |
| JV103 | Int PCR STOML tag 5' R | CTGGAACATCGTAAGGATACGC | |
| JV104 | Int PCR STOML tag 3' F | ATTATATGTGAAAATTATGGGGCATGC | |
| JV121 | Int PCR STOML tag 3' R | CTCTTTTTTTTCCACTAAAATTATTATAGG | |
| JV061 | Int PCR STOML KO 5' F | TATAGATGTGGATTATTTTACACAATTGAC | |
| NP50 | Int PCR STOML KO 5' R | CTTAATATTGATAAGTATCATGTG | |
| NP49 | Int PCR STOML KO 3' R | CCGAAAAAGTTAAAATTAATTTAC | |
| JV028 | Int PRC STOML KO 3' R | ATATTAGGCGCCTATAAATTGTACGTGTATTATTCATTACTC | |
| NP50 | Int PCR cyto-mScar 3' F | CTTAATATTGATAAGTATCATGTG | |
| NP298 | Int PCR cyto-mScar 3' R | CGTTCATGCTTTCACAAGAAC | |
| NP297 | Int PCR cyto-mScar 5' F | GCTCACCTTAAATGTTCCAC | |
| NP190 | Int PCR cyto-mScar 5' R | AGTCATATCCAGGAATAAACATAC | |

***Table S1. Primer and guide sequences for generation of repair and guide plasmids.*** *Used abbreviations: HR = homology region, F = forward primer, R = reverse primer. Overhang for restriction sites are red, restriction sites are underlined, and gRNA sequences are blue. The same primers are used for the integration PCR of cyto-mScarlet and cyto-mNG.*

**Table S2. Proteins detected in STOML-3HA-NG pulldown.**

*This table includes all proteins detected in the STOML-3HA-NG pulldown, indicated by their gene ID in the first column and the annotation in the second column. T-test significance, unique peptides, log p-value and t-test difference are indicated in the following columns. Mitochondrial prediction score is indicated in the last column as "Mito score", which refers to the ranking of the mitochondrial proteome as shown by Esveld* et al.[3]

**Table S3. Proteins detected in STOML-3HA pulldown.**

*This table includes all proteins detected in the STOML-3HA pulldown, indicated by their gene ID in the first column and the annotation in the second column. T-test significance, unique peptides, log p-value and t-test difference are indicated in the following columns. Mitochondrial prediction score is indicated in the last column as "Mito score", which refers to the ranking of the mitochondrial proteome as shown by Esveld* et al.[3]

### Movie legends

**Movie 1. 3D visualization of STOML-3HA-NG in late ring stage.** 3D visualization of STOML-3HA-NG (green) and mito-mScarlet (magenta) in ring stages with live confocal Airyscan microscopy, using Arivis 4D vision software. Fluorescent signal is segmented by manual thresholding.

**Movie 2. 3D visualization of STOML-3HA-NG in early schizont stage.** 3D visualization of STOML-3HA-NG (green) and mito-mScarlet (magenta) in early schizonts with live confocal Airyscan microscopy, using Arivis 4D vision software. Fluorescent signal is segmented by manual thresholding.

**Movie 3. 3D visualization of STOML-3HA-NG in late schizont stage.** 3D visualization of STOML-3HA-NG (green) and mito-mScarlet (magenta) in late schizonts with live confocal Airyscan microscopy, using Arivis 4D vision software. Fluorescent signal is segmented by manual thresholding.

**References**

1. Verhoef, J. M. J. *et al.* Detailing organelle division and segregation in Plasmodium falciparum. *Journal of Cell Biology* **223**, (2024).
2. Evers, F. *et al.* Composition and stage dynamics of mitochondrial complexes in Plasmodium falciparum. *Nat Commun* **12**, 3820 (2021).
3. Esveld, S. L. Van *et al.* A Prioritized and Validated Resource of Mitochondrial Proteins in Plasmodium Identifies Unique Biology. *mSphere* **6**, e00614–21 (2021).
